# Inflammatory reprogramming of human brain endothelial cells compromises blood–brain barrier integrity in Alzheimer’s disease

**DOI:** 10.1101/2025.09.26.678918

**Authors:** Rebecca L. Pinals, Md Rezaul Islam, Oisín King, Aaron Choi, Eulim Kang, Masayuki Nakano, Anjanet Tuyéras, Maeesha Tasnim Naomi, Arthur Ngo, Alan Jiang, Nhat Truong, Emre Agbas, Claudia F. Lozano Cruz, Colin Staab, Tak Ko, David A. Bennett, Alice E. Stanton, Robert Langer, Li-Huei Tsai

## Abstract

Blood-brain barrier (BBB) dysfunction is an early feature of Alzheimer’s disease (AD), yet the endothelial gene-regulatory programs involved remain incompletely understood. We integrate postmortem human single-nucleus transcriptomics with iPSC-based BBB models to define a conserved, inflammation-driven pathway that compromises barrier integrity. We identify an NF-κB-associated endothelial gene module endoM2 that is elevated in AD, inversely correlated with cognition, and enriched for inflammation and endothelial-to-mesenchymal transition signatures. Cytokine stimulation of iPSC-derived brain endothelial cells induces morphological remodeling, lipid accumulation, junctional disruption, and transcriptomic shifts that mirror endoM2. A targeted drug screen identifies the NF-κB inhibitor BAY11-7082 as protective against cytokine-induced changes. In our perfusable iPSC-derived BBB-Chip that recapitulates human BBB signatures, single-cell profiling reveals inflammatory endothelial state-specific programs reflecting those in AD brains and demonstrates that BAY11-7082 suppresses cytokine-triggered dysfunction and reverses inflammation-associated gene activation. Together, these findings position cerebrovascular inflammation as a therapeutic target to preserve BBB integrity in AD.

## Introduction

Vascular endothelial cells regulate molecular exchange between circulating blood and tissues.^1^ The blood-brain barrier (BBB) is a specialized, selectively permeable vascular interface that supports the metabolic demands of the brain by enabling local nutrient delivery and waste efflux, while restricting access of neurotoxic xenobiotics, pathogens, and endogenous compounds in circulation.^2–6^ This tightly regulated molecular transport is mediated by the BBB’s unique cytoarchitecture, comprising brain microvascular endothelial cells (BMECs) interacting closely with pericytes and astrocytes, and by BMEC-specific expression of junctional proteins and suppression of vesicular trafficking.^4,3,7,5,6^ These features result in restricted paracellular permeability and exceptionally low transcellular transport across the BBB.^2,4–6^ Though the full vascular tree of arterioles, capillaries, and venules is present in the brain, the BBB is classically defined at the level of continuous, non-fenestrated capillaries.^3,5^

Cerebral function is inextricably coupled to cerebrovascular integrity and regulation.^8,9^ The brain is one of the most highly perfused organs in the body,^10,11^ and it is estimated that no brain cell is further than ∼20 µm from a blood vessel.^12–15^ Reflecting this intimate neurovascular interdependence, BBB dysfunction is increasingly associated with dementias including Alzheimer’s disease (AD), where patients harbor focal vascular abnormalities such as vessel regression, basement membrane deposition, loss of junctional proteins, increased permeability, and neurovascular uncoupling.^5,16–21^ Genomic studies reinforce the link between vascular dysfunction and AD,^22^ with *APOE4*—the strongest genetic risk factor for AD^23–28^—being associated with accelerated BBB breakdown even in the absence of cognitive impairment^29,30^ and independent of amyloid-ß and tau pathology.^31,32^ Originating from either the CNS and periphery, ApoE4 has been shown to contribute to vascular destabilization via impaired pericyte and endothelial function.^31,33–37^ More broadly, inflammation is increasingly recognized as a key contributor to BBB breakdown in AD, again originating from both the brain parenchyma and periphery.^2,5,22,38^ Despite these emerging insights, the mechanisms by which *APOE4* and inflammation synergistically impact BMEC biology remain incompletely understood.

Recent vascular atlases have advanced our understanding of brain EC diversity by mapping transcriptional heterogeneity across regions, disease states, and individuals, profiling cohorts ranging from a few dozen to over 400 donors.^22,39–41^ Though these studies importantly catalog differentially expressed genes (DEGs), their functional consequences in disease contexts remain poorly understood. Building on these resources, we aim to define how AD impacts BBB function, both in terms of what factors the BBB is exposed to (e.g., peripheral and CNS-derived inflammation) and how BMECs respond. Here, we focus on the convergence of genetic (*APOE4*) and environmental (inflammatory) risk factors, as *APOE4* confers a systemic biological vulnerability that primes the brain for neurodegeneration in the presence of external stressors.^38^

In this study, we integrated postmortem human brain transcriptomics with iPSC-derived BMEC model systems to define AD-associated endothelial programs (**Figure 1A**). We identified an inflammation- and EndMT-enriched module, endoM2, that correlates with AD neuropathological features and cognitive decline, then modeled these signatures in BMECs and BBB-Chip systems under inflammatory stress. Pharmacologic perturbation revealed NF-κB inhibition as protective against cytokine-induced BMEC dysfunction, and single-cell profiling of the BBB-Chip connected these effects to gene signatures seen in the AD cerebrovasculature. By linking genetic and environmental AD risk factors to endothelial reprogramming, this study uncovers a possible mechanism of BBB dysfunction and highlights endothelial plasticity as a potential therapeutic target.

**Figure 1.**
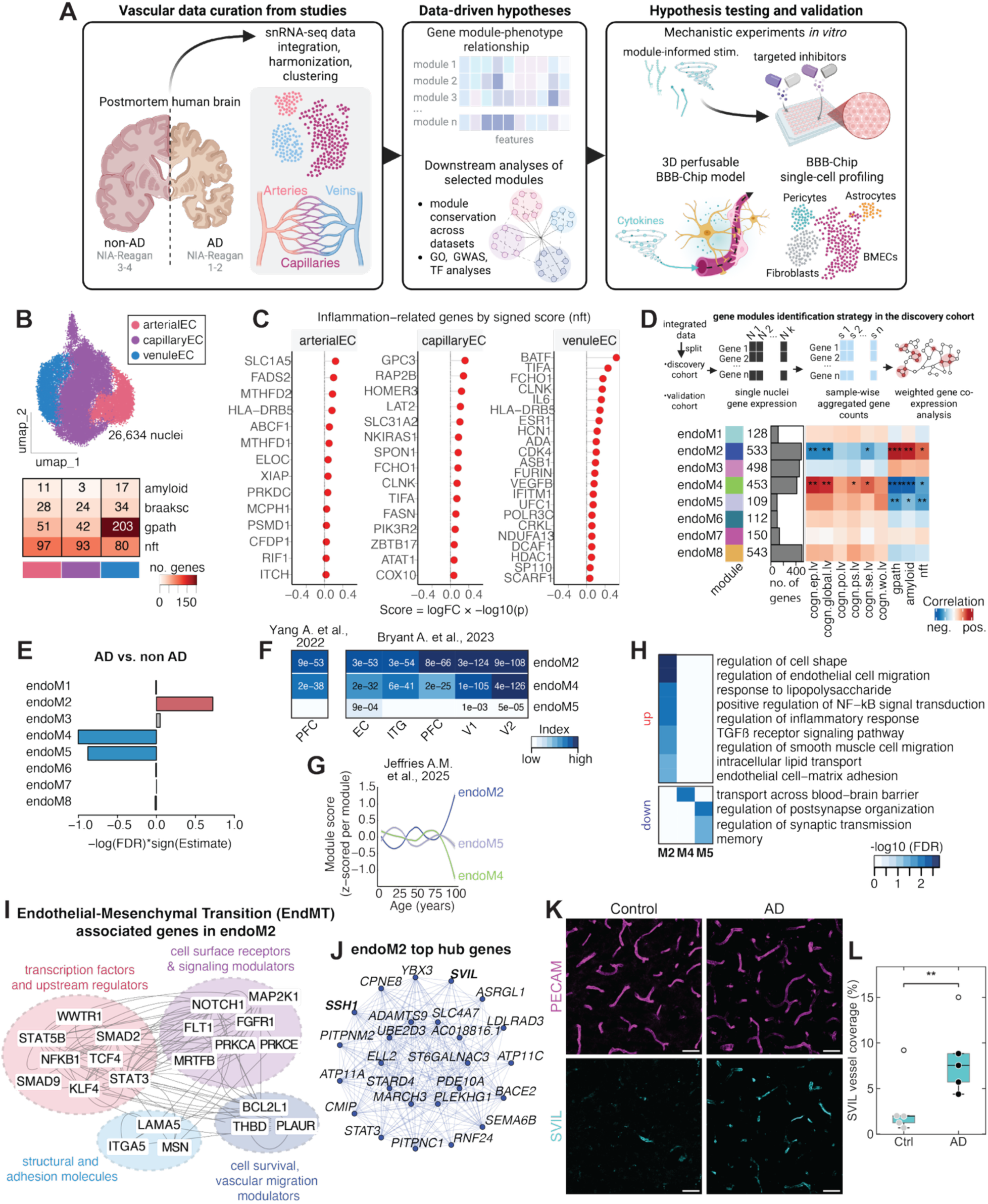
Transcriptional alterations in brain endothelial cells reveal inflammation- and migration-associated programs in Alzheimer’s disease. (A) Schematic overview of the study. Participants from ROSMAP cohorts were stratified based on the NIA-Reagan score into non-AD (NIA-Reagan: 3–4) and AD (NIA-Reagan: 1–2). Single nuclei from 431 participants from two studies were integrated and analyzed to infer data-driven hypotheses. To test the hypotheses, a perfusable human iPSCs-derived BBB model on a microfluidic chip (BBB-Chip) was employed to perform functional experiments. (B) UMAP clustering of curated endothelial cells (top) and heatmap showing number of genes significantly (FDR < 0.05) associated with AD pathology and disease progression (bottom). The color index denotes the number of genes, with darker red indicating more genes associated with a given AD phenotype. (C) Inflammatory genes significantly (FDR < 0.05), positively (red) associated with neurofibrillary tangles (nft) across endothelial cell types. (D) Schematic overview of the workflow (top) and associations between endothelial modules and AD hallmarks (bottom). Bar plots showing the number of genes in each gene module. Heatmap showing correlations between endothelial module expression and cognitive domains (cogn.epi.lv, episodic memory; cogn.global.lv, global cognition; cogn.poi.lv, perceptual orientation; cogn.ps.lv, perceptual speed; cogn.se.lv, semantic memory; cogn.wo.lv, working memory) as well as pathological measures (gpath, global pathology; amyloid, amyloid burden; nft, neurofibrillary tangles). Asterisks indicate significant associations; * *p* < 0.05, ** *p* < 0.01, *** *p* < 0.001. The color indicates Pearson’s correlation coefficient, with red representing positive and blue representing negative correlation. (E) Bar plot showing module expression in AD vs. non-AD. X-axis represents the module score [-log10(FDR)*signed(Estimate)]. Red indicates upregulation and blue represents downregulation. Data were regressed for age, sex, postmortem interval, and the percentage of mitochondrial and ribosomal reads. endoM2 is significantly upregulated in individuals with AD compared to non-AD cases. (F) Heatmap showing the conservation of AD-associated modules with endothelial differentially expressed genes (DEGs) identified in AD across datasets.^22,40^ Endothelial DEGs were curated from the indicated AD datasets. The color legend indicates Jaccard Index, with darker blue representing higher enrichment. Numbers within the boxes denote adjusted *p*-values for the modules. PFC, prefrontal cortex; EC, entorhinal cortex; ITG, inferior temporal gyrus; V1, primary visual cortex; V2, secondary visual cortex. (G) Module expression across human brains spanning the lifespan (0.4– 104 years).^52^ endoM2 module displayed a highly significant positive association with aging (x-axis) after adjusting for batch effects (adjusted *p*-value = 1.55E-20). (H) Significant biological processes associated with modules endoM2 (up), endoM4 (down), and endoM5 (down). The color scale represents -log10(FDR). (I) Protein-protein interaction network plot generated using the STRING database highlights the endoM2 module genes associated with endothelial to mesenchymal transitions (EndMT). (J) Network connectivity of top 25 hub genes in endoM2. (K) Representative images of PECAM-positive vessels (magenta) and supervillin (SVIL; cyan) immunostaining in postmortem human prefrontal cortex samples from *APOE4* carriers diagnosed with AD vs. non-AD controls. Scale bars, 50 µm. (L) Fractional coverage of supervillin on vessels as a function of AD diagnosis, analyzed for five control individuals compared to five AD patients, controlling for age and sex effects. Data points represent an individual, each the mean of two imaged regions per individual. Open markers indicate outliers identified using the interquartile range method. *p*-values were calculated using Welch’s *t*-test; ** *p* < 0.01.

## Results

### *In silico* sorting of postmortem human brain endothelial cells

To investigate endothelial cell (EC) transcriptional alterations in AD, we analyzed 26,634 EC nuclei curated from single-nucleus RNA sequencing (snRNA-seq) datasets of 431 postmortem human prefrontal cortex samples from the ROSMAP cohorts (**Figure 1A** and **Table S1**).^22,41^ Study participants were classified as AD (NIA-Reagan scores 1–2) and non-AD (scores 3–4).^42,43^ We integrated the datasets from 245 AD and 186 non-AD individuals by harmonizing data across samples and studies, then performed clustering on the EC nuclei (**Figure S1A–C**). We identified transcriptionally distinct major EC subtypes, including arterial ECs (aECs, characterized by *VEGFC* and *ALPL*), capillary ECs (cECs, characterized by *MFSD2A*, *SLC7A5*, and *INSR*), and venous ECs (vECs, characterized by *IL1R1* and *TSHZ2*) (**Figure 1B**, **Figure S1D–E**, **Table S2**), consistent with previous studies.^22,41^

### AD pathology-associated gene expression changes in endothelial cells

Next, we applied a negative binomial mixed model to identify the gene expression changes associated with disease progression and AD pathologies, while accounting for individual-level variability and other covariates including age, study batch, sex, and postmortem interval (**Table S3**). Our analysis uncovered transcripts significantly associated with amyloid burden, global pathology, and neurofibrillary tangles. Genes associated with neurofibrillary tangles were more extensively dysregulated across all EC types compared to those associated with amyloid (**Figure 1B**), underscoring the possibly broad impact of tangle pathology on endothelial gene expression and highlighting the potential role of tau-mediated vascular changes in AD.^44^ Notably, genes associated with pathologies converged on inflammatory genes (**Figure 1C**, **Figure S2**). For instance, arterial ECs displayed upregulation of *SLC1A5*, *FADS2*, *MTHFD2*, *HLA-DRB5*, *PRKDC*, *XIAP*, *ELOC*, *RIF1*, ITCH, and *MCPH1*. Capillary ECs exhibited induction of genes including *GPC3*, *RAP2B*, *HOMER3*, *NKIRAS1*, *CLNK*, *TIFA*, *FASN*, and *SLC31A2*. Venous ECs showed a particularly strong inflammatory signature, with marked upregulation of *BATF*, *TIFA*, *IL6*, *HLA-DRB5*, *ESR1*, *VEGFB*, *HDAC1*, *CDK4, CLNK*, *IFITM1*, *and SP110*, among others. Although amyloid-associated changes were more modest, they nonetheless included relevant inflammatory mediators: arterial ECs upregulated *P2RY2*, *ELOC*, and *RIPK1*, capillary ECs upregulated *NHEJ1*, and venous ECs displayed expression changes in *VEGFB*, *FKRP*, and *NCSTN* (**Figure S2A**). Interestingly, *VEGFB* expression was also associated with Braak stages (**Figure S2B**) and global pathology (**Figure S2C**). The recurrent association of *VEGFB* with various pathological measures and disease progression suggests an endothelial survival/maintenance signal in AD, consistent with its known anti-apoptotic *VEGFB* activity.^45^ Moreover, genes positively associated with global pathology also included *TNFRSF11A*, which encodes the receptor for the cytokine TNF-α, and *TICAM1*, an inducer of NF-κB activation (**Figure S2C**). Consistently, gene ontology interrogation of global pathology-associated genes revealed robust enrichment of inflammatory signaling processes, including NF-κB signaling, the JNK cascade, the MAPK cascade, and response to external stimulus (**Figure S2D**). Together, these findings suggest that endothelial cells may undergo pathology-specific transcriptional reprogramming in AD, characterized by heightened inflammatory responses alongside potential compensatory processes.

### Endothelial cell gene-regulatory network dysfunction in AD

Building on these pathology-associated increases in inflammatory processes, we next applied a network-based approach to investigate endothelial gene-regulatory programs. To unbiasedly identify such programs in AD, we constructed weighted gene co-expression networks from aggregated sample-wise transcript counts (see Methods). To this end, we divided the integrated datasets into discovery and validation cohorts (**Figure 1D**). The dataset from Mathys *et al.*, which included the endothelial nuclei from the larger number of samples (401 individuals),^46^ was designated as the discovery cohort and the dataset from Reid et al., was employed as the validation cohort.^47^ Additional validation cohorts included genes dysregulated in AD based on Yang et al. and Bryant et al.^22,40^ After regressing out phenotypic covariates including age, sex, and postmortem interval, gene co-expression network analysis identified eight endothelial modules (endoM1–endoM8), among which endoM2, endoM4, and endoM5 were significantly altered in AD (**Figure 1D–E**, **Table S4**). Expression of the endoM2 module was increased in AD, whereas endoM4 and endoM5 modules showed reduced expression (**Figure 1E**). As expected, endoM2 eigengene expression was negatively correlated with various cognitive domains and positively associated with amyloid and tau pathology, underscoring its disease relevance (**Figure 1D**). In contrast, endoM4 and endoM5 modules showed positive correlations with cognitive phenotypes and inverse associations with AD pathology, suggesting their inverse relationship with the endoM2 module. Among these modules, endoM2 and endoM4 were conserved with endothelial signatures dysregulated in AD across cortical brain regions and independent cohorts (**Figure 1F**).^48–51^

Since AD is an aging-associated disease, we asked whether the AD-associated endothelial gene modules exhibit dynamic expression changes across the lifespan. To address this, we reanalyzed another additional snRNA-seq dataset that profiled cortical samples from individuals aged 0.4 to 104 years.^52^ *In silico* sorting of 12,143 endothelial nuclei based on canonical markers (*PECAM1*, *CLDN5*, *FLT1*) revealed significant positive association of the endoM2 module with age (**Figure 1G**, **Figure S3**). Thus, we focused on the endoM2 module for further comparative analyses, based on this result and our validation cohort of the integrated datasets (**Figure S1A**) demonstrating the increased endoM2 expression in AD (**Figure S3D**). Notably, expression of the endoM2 module exhibited a dose-dependent relationship with *APOE4* risk-allele status, displaying stepwise increases across *APOE3/3*, *APOE3/4*, and *APOE4/4* genotypes (**Figure S3E**). Inspired by these observations, we stratified individuals based on their clinical consensus diagnosis of cognitive status at the time of death into three groups: no dementia, mild cognitive impairment, and dementia. We found that, relative to both no-dementia and mild cognitive impairment groups, endoM2 module expression was significantly increased in dementia (**Figure S3F**).

### Inflammatory and migratory endothelial cell phenotypes in AD

Gene ontology (GO) analyses revealed several distinct themes associated with the modules: endoM2 was associated with inflammatory processes (e.g., response to lipopolysaccharide, TGF-β, and NF-κB signaling), cell–matrix adhesion, and regulation of cell shape and migration; endoM4 with transport across the BBB; and endoM5 with synaptic function (**Figure 1H**). The endothelial migratory phenotype suggested by the endoM2 module was particularly intriguing, prompting us to ask whether these migratory phenotypes could be linked to pathological processes such as endothelial-to-mesenchymal transition (EndMT). To address this, we performed a supervised analysis and found that several genes associated with EndMT were expressed within the endoM2 module (**Figure 1I**). EndMT is a phenomenon by which ECs lose canonical characteristics, including EC markers and cell-cell junctions, delaminate from the vessel wall, and acquire a motile, mesenchymal phenotype; a transition classically described in development, tumor invasion and metastasis, and cardiovascular disease.^53–57^ The EndMT genes in endoM2 include transcriptional drivers (e.g., *NFKB1*, *SMAD2*, SMAD9, *STAT3*, *STAT5B*, *TCF4*, *KLF4*), signaling mediators of inflammation (e.g., *FLT1*, *FGFR1*, *NOTCH1*, *MAP2K1*), structural and cytoskeletal components (e.g., *MSN*, *ITGA5*, *LAMA5*), and regulators of vascular migration (e.g., *PLAUR*, *BCL2L1*, *THBD*) (**Figure 1J**), suggesting a link between inflammatory signaling and EndMT in AD ECs. In line with this, we identified several EMT-related transcription factors (e.g., *ETS1*, *ETS2*, *SNAI1*) and inflammatory signaling regulators (e.g., *NFIL3*, *BHLHE40*, *NR4A1*) as potential upstream drivers of the endoM2 module (**Figure S4A**). Furthermore, analyses of AD-risk-associated genes within the endoM2 module highlighted transcriptional regulators such as *STAT1*, *STAT2*, *IRF2*, *RELB*, and *ETS1*, involved in NF-κB signaling and mesenchymal transition programs (**Figure S4B**).

Network connectivity analysis identified *SVIL* and *SSH1* as top hub genes within the endoM2 module (**Figure 1J**). *SVIL* encodes supervillin, a membrane-associated protein that links the actin cytoskeleton to the plasma membrane and regulates cell shape and motility.^58,59^ *SSH1* encodes slingshot protein phosphatase 1 (SSH1), a key activator of cofilin that drives actin filament turnover and cytoskeletal remodeling, thereby enhancing cell migration.^60^ Both *SVIL* and *SSH1* have been implicated in EndMT processes, particularly in the context of cancer,^61–66^ and their presence as hub genes within the upregulated endoM2 module further supports a shift toward an EndMT-like state in AD capillary ECs. Notably, supervillin fractional coverage on brain vessels was significantly elevated in AD brains from the MAP cohort compared to age-matched controls (all *APOE4* carriers), increasing from 3.02% to 8.29% average fractional coverage based on immunostaining (**Figure 1K–L**). Moreover, vessel coverage of supervillin (*p* = 0.026) and SSH1 (*p* = 0.021) were both significantly correlated with brain amyloid load (**Figure S5**). These observations support a model in which AD-associated inflammation may drive *SVIL*/*SSH1*-mediated cytoskeletal reprogramming in brain endothelium.

### Module-to-phenotype connection in iPSC-derived BMECs

We aimed to phenotypically reproduce and mechanistically interrogate endoM2 transcriptional changes *in vitro* using human induced pluripotent stem cells (iPSCs) to differentiate BMECs (iBMECs; see Methods). iBMECs were validated to possess canonical endothelial markers and lack epithelial markers (**Figure S6A**), consistent with prior transcriptomic analyses.^67^ Flow cytometry further confirmed capillary EC identity, demonstrated by co-expression of CD31 and CD34 (**Figure S6B**). We differentiated iBMECs from an isogenic pair of iPSC lines: homozygous *APOE3* and CRISPR-edited homozygous *APOE4*. At baseline, *APOE4* iBMECs clustered separately from *APOE3* iBMECs by principal component analysis (**Figure S6C**) and displayed transcriptomic features associated with endoM2, including GO terms related to cell migration and inflammation (**Figure S6D**).

To determine relevant stimulation conditions that connect to the endoM2 module observed in postmortem human BMECs, we noted several cytokine-related pathways including IL-1 and IL-6 signaling were specifically enriched in endoM2 (**Figure S3C**). IL-1, IL-6, and TNF-α have been reported to be dysregulated in neurodegenerative diseases.^68–70^ Consistent with this, supervised analyses of the endoM2 module revealed several receptors (*TNFRSF10B*, *TNFRSF10D*) and proteins (*TNFAIP1*, *TRAF3IP2*) related to TNF signaling (**Table S4**). Additional candidate stimuli were informed by biological terms enriched in endoM2, including response to lipopolysaccharide (LPS) and lipid-related processes (**Figure 1H**). Based on these observations, our stimulation panel included (i) LPS, (ii) oleic acid to trigger lipid accumulation, and (iii) a pro-inflammatory cytokine cocktail (TNF-α, IL-1β, and IL-6) (**Figure 2A**). Although we observed phenotypic responses in both *APOE3* and *APOE4* iBMECs, we focused on *APOE4* due to their more pronounced transcriptional shifts with AD pathology in postmortem human brain tissue (**Figure S3E**). Each stimulus was applied to iBMECs for 24 hours, followed by live staining with Hoechst to visualize nuclei, BODIPY neutral lipid stain for lipid droplets, or pHrodo-LDL for lipoprotein particle uptake (see Methods). To monitor tight junction dynamics in live cells, we used an iPSC line expressing the tight junction protein ZO-1 fused to monomeric enhanced green fluorescent protein (mEGFP).

**Figure 2.**
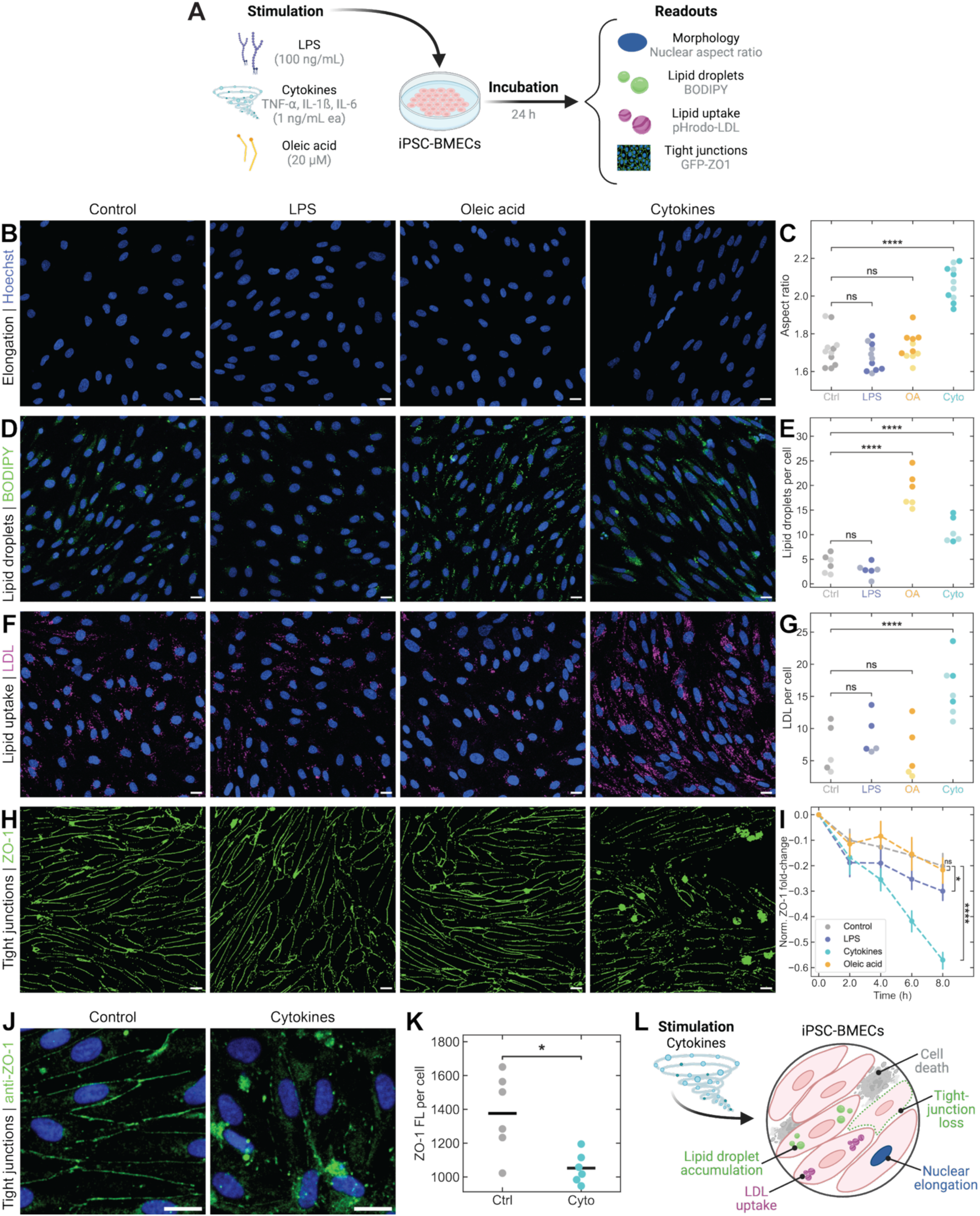
iPSC-derived BMECs exhibit endoM2-related phenotypes under inflammatory stress. (A) Schematic of iBMECs exposed to stimulation conditions informed by gene ontology analysis of the endoM2 module: lipopolysaccharide (LPS, 100 ng/mL), a cytokine cocktail (TNF-α, IL-1β, and IL-6, 1 ng/mL each), or oleic acid (OA, 20 µM) for 24 h. Representative live-cell images and corresponding quantification of iBMECs responses: (B-C) nuclear morphology (Hoechst, blue), (D-E) lipid droplets (BODIPY, green), and (F-G) LDL uptake (pHrodo-LDL, magenta) all in *APOE4* iBMECs; and (H-I) ZO-1 tight junctions (ZO-1-mEGFP, green) in *ZO-1*-mEGFP iBMECs. Scale bars, 20 µm. Data points represent biological replicates, each the mean of *n* ≥ 2 imaged regions per well. Independent iBMEC batches are indicated by light and dark marker colors. (C, E, G) Statistical testing was performed using a linear mixed-effects model with sample type as a fixed effect and batch as a random intercept. *p*-values were calculated using Wald tests of the fixed-effect coefficients. (I) Error bars represent 95% confidence intervals of the mean. *p*-values were calculated using Welch’s *t*-test comparing each treatment to control at the final timepoint. (J) Immunostaining of ZO-1 in *APOE4* iBMECs under control and cytokine-treated conditions (ZO-1, green; Hoechst-stained nuclei, blue). Scale bars, 20 µm. (K) Quantification of ZO-1 along cell junctions normalized to nuclei number. Data points represent biological replicates, each the mean of two imaged regions per well, and horizontal lines represent the group mean. *p*-values were calculated using Welch’s t-test; **** *p* < 0.0001, *** *p* < 0.001, ** *p* < 0.01, * *p* < 0.05, ns *p* > 0.05. (L) Summary schematic of effects imparted on endothelial cells from cytokine stimulation.

Of the tested stimulation conditions, cytokine treatment uniquely elicited multiple phenotypic changes in iBMECs: cellular elongation (quantified by increased nuclear aspect ratio; **Figure 2B–C**), elevated lipid droplet load (approaching the oleic acid positive control; **Figure 2D–E**), increased LDL uptake (**Figure 2F–G**), and disruption of ZO-1 tight junctions (diminished mEGFP fluorescence at cell borders; **Figure 2H–I**). Because the *ZO-1*-mEGFP-iBMECs were not in *APOE4* background, we further confirmed these findings in fixed *APOE4*-iBMECs immunostained for ZO-1, similarly demonstrating tight junction disruption via decreased ZO-1 presence along cell borders (**Figure 2J–K**). These changes parallel the shift from a homeostatic BBB phenotype toward the disease-associated state suggested by the postmortem human snRNA-seq data. Mechanistically, these morphological and junctional alterations are consistent with prior work showing that pro-inflammatory cytokines destabilize tight junctions by promoting F-actin reorganization into stress fibers.^2,71^ The spindle-like nuclear shape mirrors mesenchymal cells such as fibroblasts,^72^ while increased lipid droplet content and elevated LDL uptake are atypical for healthy microvascular ECs and suggest a shift toward a pro-inflammatory, metabolically altered state.^73,74^ ZO-1, a cytoplasmic adaptor linking transmembrane adhesion complexes to the actin cytoskeleton,^5^ likely undergoes disassembly and mislocalization under cytokine exposure, with the presence of mEGFP aggregates due to vesicular trafficking or extracellular shedding. In parallel, ICAM-1 expression increased following cytokine treatment as expected (**Figure S7**).^75^ Taken together, these results support a model in which cytokine exposure compromises iBMECs through cytoskeletal remodeling, lipid accumulation, and redistribution of junctional proteins away from cell borders (**Figure 2L**), reflecting key features of EndMT-like remodeling.

### Cytokines drive iBMECs toward AD-associated transcriptional states

To transcriptionally quantify whether the AD-associated endothelial program identified in the postmortem brain can be modeled under inflammatory conditions *in vitro*, we treated iPSC-derived BMECs from isogenic *APOE3* and *APOE4* lines (primary cohort) with the same cytokine cocktail (TNF-α, IL-1β, and IL-6) and performed bulk RNA sequencing (**Figure 3A**). *APOE4* iBMECs exposed to cytokines yielded 3,211 up-regulated and 3,138 down-regulated genes (**Figure 3B**). Differential expression analysis revealed robust transcriptional activation in response to cytokine stimulation in both genotypes, with a more pronounced and stronger response in *APOE4* iBMECs (**Figure S8A**). We validated this observation in the primary cohort (*APOE3/3* parental) by repeating the experiment in another iPSC isogenic pair (*APOE4/4* parental) from which iBMECs were differentiated (replication cohort), and observed a high correlation of the common DEGs between the two cell lines (**Figure S8B–C**). We focused on the primary cohort for downstream analyses and the upregulated genes included canonical inflammation-responsive transcripts (e.g., *CXCL8*, *NFKB1*) and stress-associated markers (e.g., *SSH1*) in both *APOE3* and *APOE4* backgrounds.

**Figure 3.**
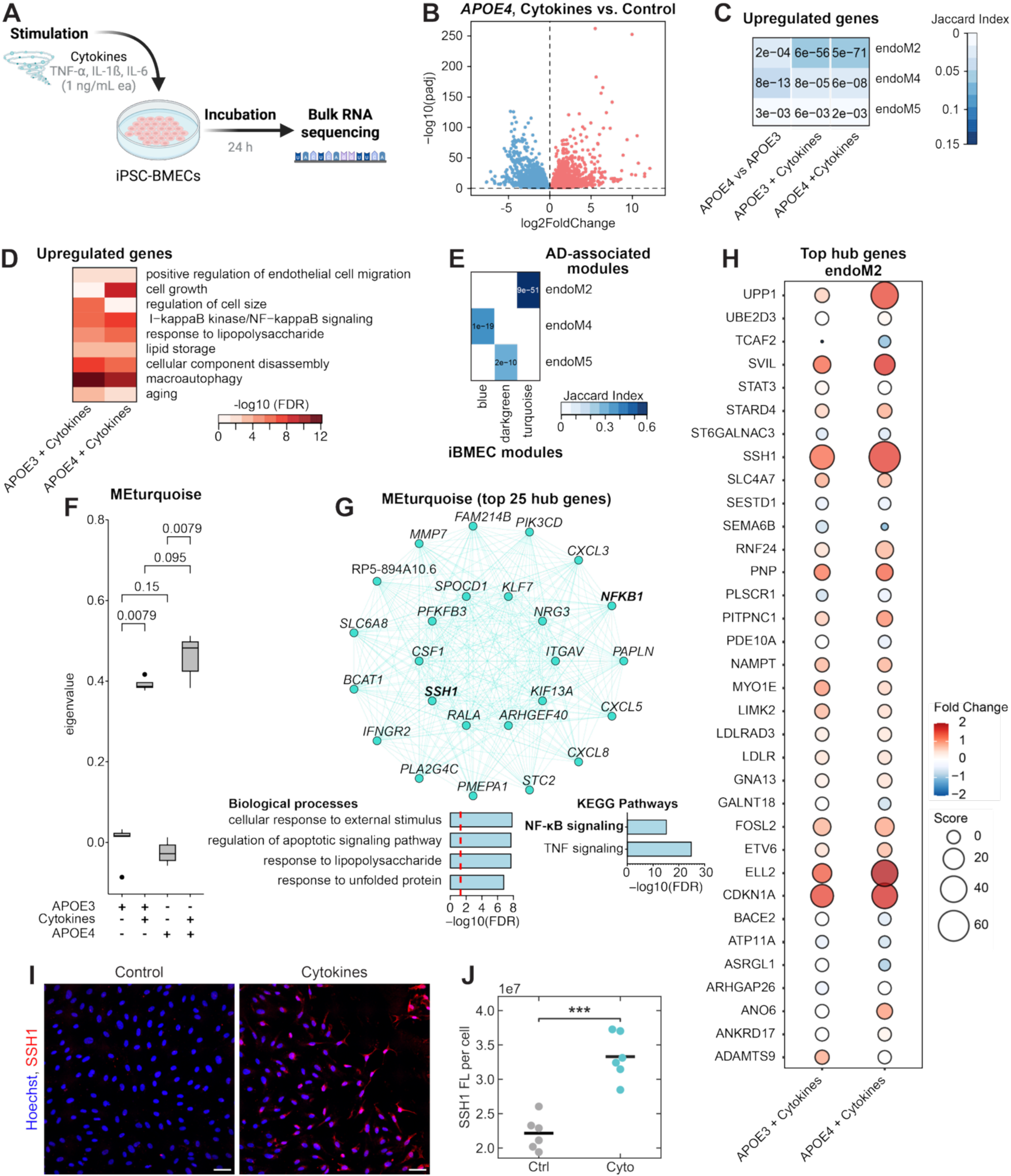
Cytokine stimulation of iBMECs aligns with AD-associated transcriptional programs. (A) Schematic of iBMEC cytokine stimulation followed by bulk RNA-seq. (B) Volcano plot showing significantly (FDR < 0.05) upregulated (red) and downregulated (blue) genes in *APOE4* iBMECs after cytokine treatment. (C) Gene-overlap analysis of cytokine-upregulated DEGs with AD-associated endothelial modules (endoM2, endoM4, endoM5) from postmortem snRNA-seq data. Color indicates the Jaccard Index. Numbers indicate adjusted *p*-values. (D) GO enrichment of the upregulated genes following the cytokine cocktail treatment in *APOE3* and *APOE4* iBMECs. Color indicates significance. (E) Weighted gene co-expression network analysis (WGCNA) of cytokine-treated iBMECs identifying the turquoise iBMEC module with the strongest overlap with AD-associated endoM2. Color indicates the Jaccard Index. (F) Turquoise iBMEC module co-expression across treatment conditions in iBMECs. Numbers indicate *p*-values. Significance was tested with the Kruskal–Wallis test. In the boxplots, the middle line marks the median value. The top and bottom edges of the box show the 75th and 25th percentiles, respectively. The whiskers extend to the minimum and maximum values that fall within 1.5 times the interquartile range. (G) (Top) Top 25 hub genes of the turquoise module, including *NFKB1*, *SSH1* (bolded). (Bottom) Gene ontology biological process (left) and KEGG pathways (right) enrichment of the turquoise module genes highlighting their roles in increased inflammatory-associated processes and NF-κB/TNF signaling pathways, respectively. Dashed red lines indicate the significance cutoff. (H) Bubble plots showing differentially expressed hub genes of endoM2 in cytokine-treated *APOE3* and *APOE4* iBMECs. Color index indicates fold change and dots representing the gene score [-log10(FDR)* fold change]. Notably, both *SSH1* and *SVIL* expression increased in cytokine-treated *APOE3* and *APOE4* iBMECs, with relatively higher levels observed in *APOE4* iBMECs. (I) Immunostaining and (J) quantification of SSH1 in *APOE4* iBMECs under control and cytokine-treated conditions (SSH1, red; Hoechst-stained nuclei, blue; scale bars, 50 µm). Data points represent biological replicates, each the mean of two imaged regions per well, and horizontal lines represent the group mean. *p*-values were calculated using Welch’s *t*-test; *** *p* < 0.001.

We compared the differentially expressed genes (DEGs) in cytokine-exposed iBMECs (see **Table S5–S6** for full DEG list) with AD endothelial modules endoM2, endoM4, and endoM5 from the postmortem snRNA-seq data. Jaccard analysis indicated significant overlap between cytokine-upregulated genes and AD-associated endothelial modules (**Figure 3C**). This overlap was consistent across *APOE3* and *APOE4* backgrounds, although the significance of overlap (**Figure 3C**), particularly for endoM2, and the number of DEGs (**Figure S8A**) were relatively higher in *APOE4* iBMECs treated with cytokines, perhaps indicating a genetic-environment interaction contributing to endothelial vulnerability. GO analysis of the upregulated genes from both genetic backgrounds revealed enrichment for endothelial migration, NF-κB signaling, lipopolysaccharide response, lipid storage, and aging-related processes (**Figure 3D**). These results indicate that inflammatory cytokines are sufficient to promote a substantial portion of the AD-associated endothelial transcriptional changes observed *in vivo*, highlighting inflammation as a central upstream regulator of disease-associated endothelial vulnerability and dysfunction.

To contextualize these findings within endothelial gene-regulatory networks, we employed a weighted gene co-expression network analysis to the cytokine-treated iBMEC transcriptomes from the primary cohort, identifying 27 gene co-expression modules (iBMEC modules) (see Methods). Among these, we focused on three modules that showed significant overlap with human AD-associated gene modules (*see* Figure 1, **Figure 3E**). Interestingly, the turquoise iBMEC module, which includes genes such as *SSH1*, *SVIL*, and *NFKB1*, showed the highest overlap with the postmortem human brain-derived endoM2 module (**Figure 3E**, **Table S7**) and was upregulated in cytokine-treated *APOE3* and *APOE4* ECs (**Figure 3F**). GO enrichment of the turquoise iBMEC module revealed upregulation of processes involved in response to external stimulus, unfolded protein, lipopolysaccharide, and apoptotic signaling, and pathways related to NF-κB and TNF signaling, all consistent with inflammation-driven endothelial stress (**Figure 3G**). Top brain-derived endoM2 hub genes differentially expressed by cytokine treatment (**Figure 3H, Figure S8D**) further suggest that cytokine stimulation drove an EndMT-like transcriptional program, which is relatively more pronounced in *APOE4* iBMECs than APOE3 iBMECs. Canonical EndMT drivers (*SNAI1*, *ZEB2*) and associated regulators (*SVIL*, *SSH1*, *LGALS3*) were significantly upregulated in *APOE4* iBMECs, alongside fibroblast-associated genes (*TAGLN*, *FN1*, *TNC*), indicating a shift away from endothelial homeostasis (**Table S6**). Staining confirmed increased SSH1 in cytokine-treated *APOE4* iBMECs relative to untreated control (**Figure 3I–J**). In parallel, lipid metabolism and storage genes (*PLIN2*, *DGAT2*) and the LDL receptor (*LDLR*) were elevated in *APOE4* iBMECs (**Table S6**), in line with the observed increase in lipid droplet accumulation and LDL uptake (**Figure 2D–G**). Cytokine treatment also triggered strong induction of inflammatory transcripts (*ICAM1*, *VCAM1*, *TLR4*, *CXCL12*, *EFNB1*, *CASP4*) in *APOE4* iBMECs, suggesting activation of a pro-inflammatory endothelial state. Conversely, several junctional and barrier-stabilizing genes were downregulated (*PECAM1*, *TJP2*, *JAM3*, *ESAM*), while *JAM2* was upregulated (**Table S6**), consistent with impaired intercellular adhesion and loss of barrier integrity in *APOE4* iBMECs. Together, these changes demonstrate that cytokine exposure recapitulates the transcriptional hallmarks of EndMT, metabolic stress, and inflammation observed in AD-associated endothelium *in vivo*.

### Targeted drug panel identifies NF-κB inhibition as protective in iBMECs

Based on the cytokine-driven alterations in iBMECs observed at both the phenotypic and transcriptomic levels, we tested whether these effects could be attenuated by pre-treatment with small-molecule inhibitors chosen to target pathways linked to *SVIL* and *SSH1* (**Figure 4A**). Notably, both *SVIL* and *SSH1* are members of endoM2 (**Figure 1J, Table S4**) and the turquoise iBMEC modules (**Table S7**), are increased in cytokine-treated iBMECs (**Figure 3H, Figure S8D**), and are linked to EndMT regulation. We assembled a focused panel of six compounds informed by prior literature, including both pharmacological agents and natural products: BAY11-7082^76^ (BAY11) and resatorvid^76^ (both associated with *SVIL*-related inflammation), and sennoside A,^77^ hypericin,^77^ Wnt antagonist III (Box5),^78^ and Dvl-PDZ domain inhibitor II^78^ (associated with *SSH1*-related cytoskeletal changes and Wnt signaling). While selected based on these gene connections, the compounds also exert broader effects: BAY11-7082 as a general NF-κB pathway inhibitor,^79,80^ resatorvid as a TLR4 antagonist,^81,82^ sennoside A as possessing broad anti-inflammatory properties,^83^ hypericin as a PKC inhibitor,^84^ and Box5^85^ and Dvl-PDZ domain inhibitor II^86^ as Wnt pathway inhibitors.

**Figure 4.**
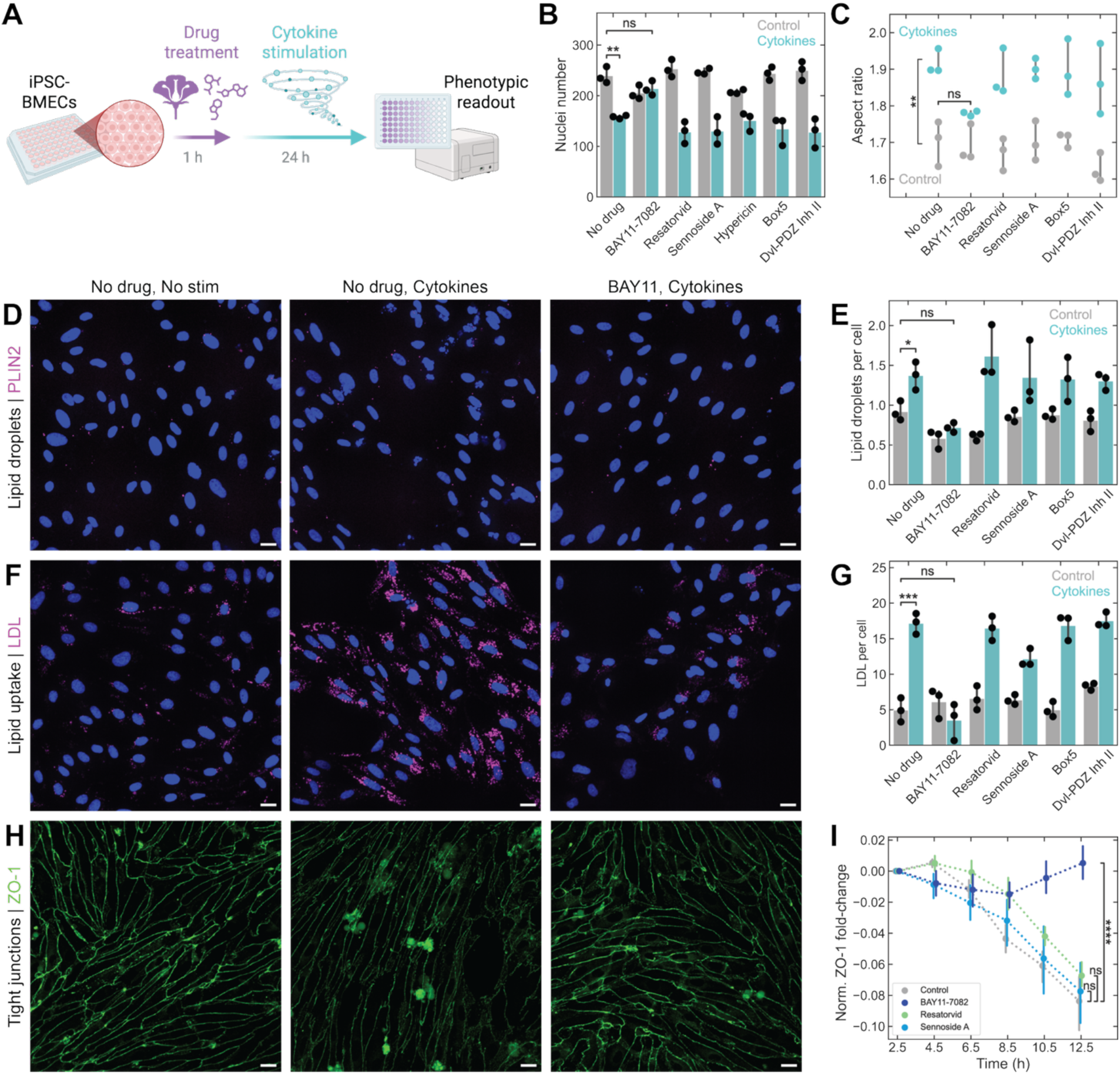
BAY11-7082 blocks cytokine-induced changes in iBMECs. (A) Schematic of iBMEC drug pre-treatment and cytokine stimulation followed by high-throughput live imaging (see Methods for details on drug panel; cytokine cocktail: TNF-α, IL-1β, and IL-6, 1 ng/mL each, 24 h). Quantification of nuclei in *APOE4* iBMECs showing (B) total nuclei and (C) nuclear morphology. Representative images and quantification of (D-E) lipid droplets in *PLIN2*-RFP-*APOE4* iBMECs (PLIN2, magenta), (F-G) LDL uptake in *APOE4* iBMECs (pHrodo-LDL, magenta), and (H-I) ZO-1 tight junctions in *ZO-1*-mEGFP iBMECs (ZO-1, green). Scale bars, 20 µm. Data points represent biological replicates, each the mean of *n* ≥ 2 imaged regions per well. Error bars represent 95% confidence intervals of the mean. *p*-values were calculated using Welch’s *t*-test; **** *p* < 0.0001, *** *p* < 0.001, ** *p* < 0.01, * *p* < 0.05, ns *p* > 0.05.

To test these compounds, we generated an *APOE4* iPSC line in which perilipin-2 (*PLIN2*), a lipid droplet-associated protein, was tagged with red fluorescent protein (RFP) (**Figure S9**). Among the compounds tested, we found that BAY11 prevented cytokine-induced cell death (**Figure 4B**) and morphological changes, reducing the nuclear aspect ratio to that of unstimulated controls (**Figure 4C**). BAY11 further suppressed lipid droplet accumulation (**Figure 4D–E**) and LDL uptake (**Figure 4F–G**), while preserving ZO-1 tight junction integrity (**Figure 4H–I**). Tested compounds alone did not cause any significant differences in comparison to the untreated control (**Figure 4**). Because hypericin showed no rescue of cellular elongation and possesses photosensitizing properties and overlapping fluorescence, this compound was thus excluded from downstream tests.^87^

### An iPSC-derived BBB-Chip with vascular flow emulates the human BBB *in vitro*

Having established the connection between human postmortem brain transcriptomics to an *in vitro* model, we next extended these findings to a more physiologic, multicellular BBB context. We built a BBB-Chip composed of endothelial cells, astrocytes, and pericytes. These vascular cell types were differentiated separately from iPSCs and, together with human fibroblasts, co-encapsulated in a 3D hydrogel and seeded into a microfluidic device (**Figure 5A**).^88^ Over approximately a week in co-culture, cells self-assembled into 3D microvascular networks, with pericytes and astrocytes homing to the microvasculature (**Figure 5B**). Importantly, this modular platform enables precise control over cell composition and the ability to employ cell-type–specific live readouts. Accordingly, iBMECs were differentiated from the *PLIN2*-RFP-*APOE4* iPSC line to selectively visualize endothelial lipid droplet accumulation, or from the *ZO-1*-mEGFP iPSC line to report on tight-junction dynamics, while astrocyte and pericyte support cells were derived from *APOE3* iPSCs. We modeled cytokine-driven barrier disruption in the BBB-Chip under flow, and tested whether BAY11 mitigates these alterations **(Figure 5C**). Vessels formed hollow lumens in the BBB-Chips that could be perfused to mimic cerebrovascular blood flow (**Figure 5D**), thus allowing us to model peripheral inflammatory insult impacting the BBB.

**Figure 5.**
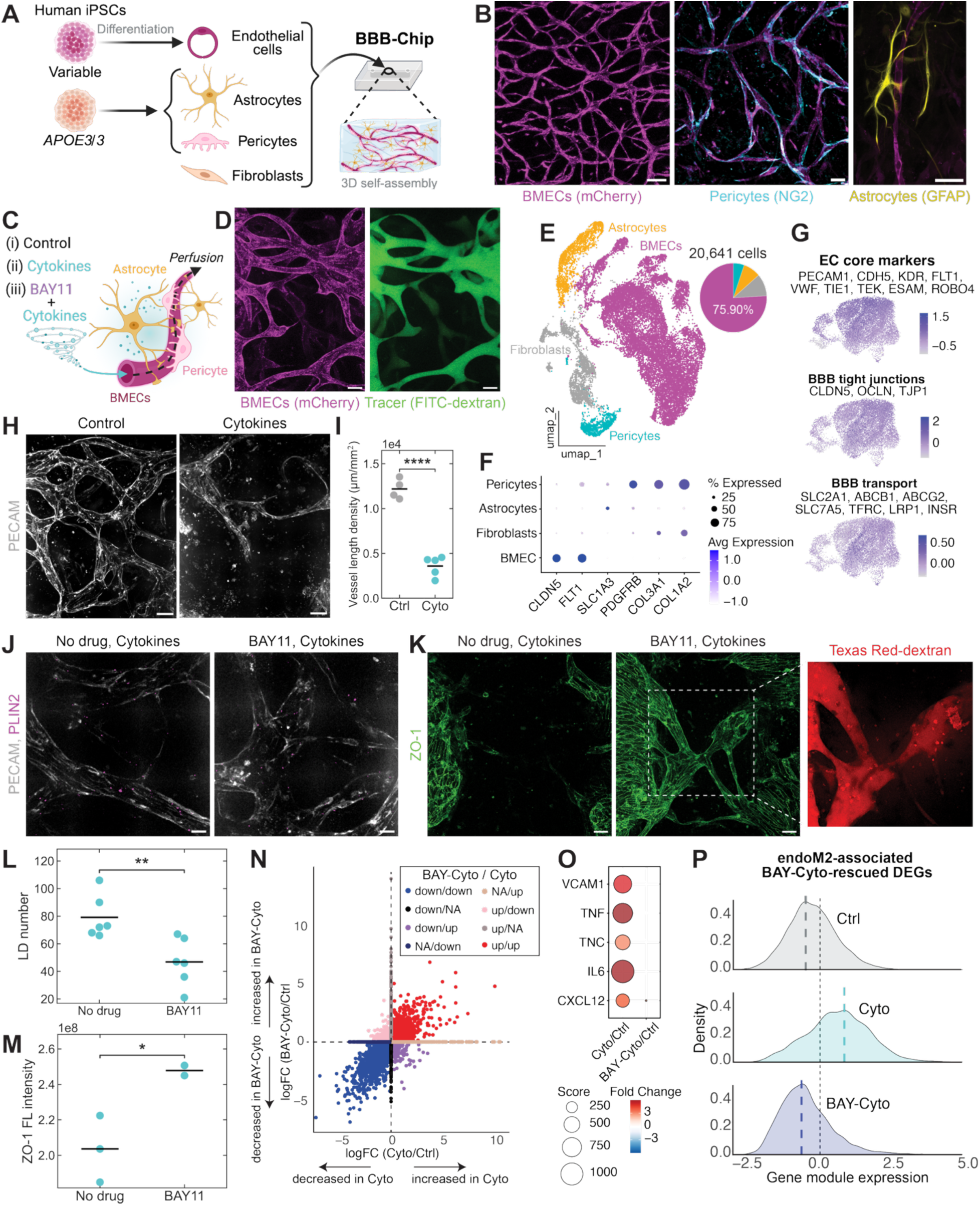
iPSC-derived BBB-Chip as a platform to model cytokine-induced changes *in vitro*. (A) Schematic of the BBB-Chip model composed of iPSC-derived BMECs, astrocytes, and pericytes, combined with human fibroblasts and co-encapsulated in a 3D hydrogel within a microfluidic chip device. (B) Representative images of BBB-Chips containing iBMECs that form microvascular networks (mCherry, magenta; scale bar, 100 µm), with pericytes (NG2, cyan; scale bar, 50 µm) and astrocytes (GFAP, yellow; scale bar, 50 µm) homing to the vasculature. (C) Schematic of BBB-Chip treatment conditions applied by perfusion through microvascular networks: control, cytokine stimulation (TNF-α, IL-1β, and IL-6, 1 ng/mL each, 18 h), and BAY11 pre-treatment (5 µM, 1 h) followed by cytokine stimulation. (D) Representative images of BBB-Chip with tracer (FITC-dextran 70 kDa, green) flowing through iBMEC vessels (mCherry, magenta; scale bars, 50 µm). (E) UMAP plot of cell types identified in the scRNA-seq from BBB-Chip. A total of 20,641 single cells were analyzed from a total of 16 samples (Ctrl: n = 5; Cyto: n = 5, BAY-Cyto: n = 6). (F) Dot plots showing cell type–specific marker gene expression, with dot size indicating the percentage of cells expressing each gene and color representing the average scaled expression. (G) Expression of core EC markers, BBB tight junctions and transport module genes expression in iBMECs. Color code represents average expression. (H) Representative images of control and cytokine-exposed BBB-Chips (PECAM, gray; scale bars, 50 µm). (I) Corresponding vessel length density (total centerline length per ROI area). (J) Representative images of BBB-Chips with *PLIN2-*RFP-*APOE4* iBMECs exposed to cytokines with or without BAY11-7082 pre-treatment (PLIN2, magenta; PECAM, gray; scale bars, 20 µm). (K) Representative images of BBB-Chips with *ZO-1*-mEGFP iBMECs exposed to cytokines with or without BAY11-7082 pre-treatment (ZO-1, green; scale bars, 50 µm). Inset demonstrates perfusion of tracer through BAY11-treated BBB-Chip (Texas Red-dextran, red). (L) Quantification of lipid droplets in endothelial cells. (M) Quantification of ZO-1 fluorescence (FL) intensity. (I, L, M) Data points represent biological replicates, each the mean of *n* ≥ 2 technical replicates in (L). *p*-values were calculated using Welch’s *t*-test; *** *p* < 0.001, ** *p* < 0.01, * *p* < 0.05, ns *p* > 0.05. (N) Comparative DEGs analyses among treatment conditions (Ctrl: control; Cyto: cytokine-treated; BAY-Cyto: BAY11-7082 pretreatment plus cytokine-treated). The x-axis shows fold changes of DEGs (FDR < 0.05) in Cyto vs Ctrl, while the y-axis shows the fold changes of DEGs in BAY-Cyto vs Ctrl. We focused on the subset of DEGs shown in purple for downstream analyses. (O) Dot plots showing expression of key inflammatory genes in Cyto/Ctrl and BayCyto/Ctrl groups. Color index indicates fold change and dots represent the gene score [-log10(FDR)* fold change]. (P) Density plots showing expression of DEGs in endoM2 module across treatment conditions. Dashed bold vertical lines indicate the peak density of gene module expression for each condition. Significance was tested with the Kruskal-Wallis test followed by Dunn’s multiple comparisons test. Cyto vs Ctrl: increased expression, adjusted *p*-value < 0.0001, BAY-Cyto vs Cyto: decreased expression, adjusted *p*-value < 0.0001.

First, we performed single-cell RNA-seq of BBB-Chips to evaluate the similarity of iBMECs to endothelial cells from the human brain. We analyzed 20,641 cells classified as endothelial cells (*PECAM1*, *CLDN5*, *FLT1*), pericytes (*PDGFRB*), astrocytes (*SLC1A3*), and fibroblasts (*COL3A1*, *COL1A2*), with endothelial cells comprising ∼76% of the dataset (**Figure 5E–F**). Endothelial cells expressed canonical EC identity markers (*PECAM1*, *CDH5*, *KDR*, *FLT1*, *VWF, TIE1*, *ESAM*, *ROBO4*), as well as BBB tight-junction genes (*CLDN5*, *OCLN*, *TJP1*) and transporters (*SLC2A1*, *ABCB1*, *ABCG2*, *SLC7A5*, *TFRC*, *LRP1*, *INSR*), supporting their physiological relevance in the BBB model (**Figure 5G**). Comparison with reference datasets further confirmed that iPSC-derived ECs in the BBB-Chip correspond primarily to brain capillary ECs (**Figure S10**), expressing both pan-endothelial markers and human brain–enriched EC markers. Together, these results demonstrate that the BBB-Chip recapitulates the *in vivo* endothelial gene expression program and provides a tractable platform to dissect context-dependent transcriptional changes.

### BAY11 prevents inflammatory barrier disruption in the BBB-Chip

We next investigated phenotypic and transcriptomic changes imparted by cytokines in the BBB-Chip under flow. Under cytokine stimulation, iBMEC vessels exhibited pronounced vascular retraction and network destabilization (**Figure 5H–I**). BAY11 attenuated multiple cytokine-induced phenotypes, reducing endothelial lipid droplets (**Figure 5J, L**) and preserving ZO-1–positive junctions (**Figure 5K, M**). From scRNA-seq analysis of the BBB-Chip, differential expression analysis identified a subset of DEGs that were upregulated in cytokine-stimulated iBMECs but attenuated by BAY11 pre-treatment (**Figure 5N**, **Table S8–9**). These DEGs were selected for downstream analyses. Notably, inflammatory and adhesion-related genes, including *VCAM1*, *TNF*, *TNC*, *IL6*, and *CXCL12*, were strongly upregulated by cytokine exposure and significantly reduced following BAY11 treatment (**Figure 5O**). A similar pattern was observed for a set of lipid-associated genes (*LPL*, *LIPG*, *ANGPTL4*, *ACOT8*, *SQLE*, *INSIG1*) (**Figure S11**). Moreover, cytokine-induced but BAY11-rescued transcriptional program included endoM2 module genes, such as transcription factors involved in cellular apoptosis (*GLIS3*,^89^ *EBF4*^90^) inflammatory mediators (*NFKB1*, *NFKBIZ*, *IRAK3*, *IL3RA*, *IL4R*), immune response regulators (*IRF2*, *SRSF7*, *PPP4R2*), and factors for cytoskeletal reorganization and adhesion remodeling consistent with their roles in EMT (*SYNJ2*,^91^ *ETS1*,^92^ *CRIM1*,^93^ *MAP3K13*^94^). Collectively, these endoM2-associated DEGs represent a cytokine-driven program that is specifically rescued by targeting NF-κB, as reflected by a composite score showing increased expression following cytokine stimulation and reduction with BAY11 pre-treatment (**Figure 5P**).

### Single-cell profiling of the BBB-Chip reveals iBMEC inflammatory state-specific gene signatures conserved in AD brains

Finally, we investigated how inflammatory stimulation affects the potential transcriptional states of iBMECs in the BBB-Chip (**Figure 6A**). Clustering of 15,667 endothelial single cells from the BBB-Chip revealed ten transcriptionally distinct subclusters (Endo-1 to Endo-10), each characterized by unique biological processes such as oxidative phosphorylation, WNT signaling, VEGF signaling, DNA damage repair and response to misfolded proteins (**Figure 6B**). Notably, three subclusters (Endo-4-6) enriched for NF-κB signaling, TGFβ signaling, and intrinsic apoptotic signaling were broadly classified as the inflammatory endothelial state (**Figure 6B, Figure S12A**). Projection of cells by treatment condition further demonstrated that cytokine stimulation shifted endothelial cells from a homeostatic to an inflammatory state (**Figure 6C**).

**Figure 6.**
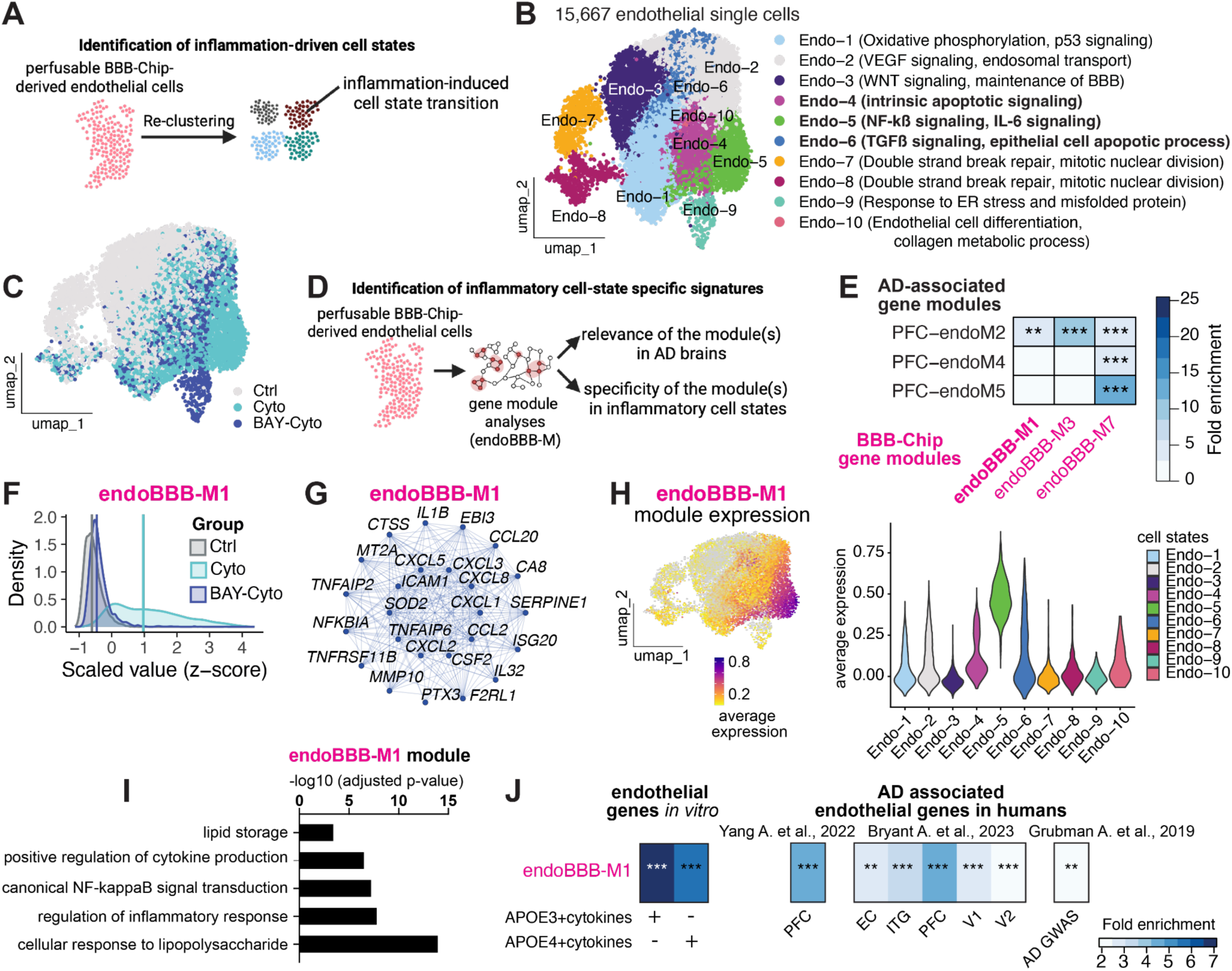
Identification of inflammation-driven endothelial cell state(s) and inflammatory state–specific gene signatures using the BBB-Chip model. (A) Schematic overview of re-clustering endothelial cells from the perfusable BBB-Chip to identify inflammation-induced endothelial cell states. (B) UMAP embedding of 15,667 endothelial single cells from the BBB-Chip model identifying ten distinct endothelial cell states (Endo-1 to Endo-10) with associated biological pathways such as maintenance of BBB, endothelial differentiation, oxidative phosphorylation, VEGF signaling, DNA repair, inflammatory and apoptotic pathways (NF-κB/IL-6 signaling, TGFβ signaling), and potential compensatory responses to endoplasmic reticulum stress and misfolded protein. We define the Endo-4, Endo-5 and Endo-6 clusters collectively as the inflammatory endothelial state (**Figure S12A**), which is enriched with apoptotic signaling, TGFß signaling and NF-κB signaling. (C) UMAP visualization of endothelial cell states under different treatment conditions: control (Ctrl, gray), pro-inflammatory cytokines-treated (Cyto, teal), and BAY11-7082 plus cytokines-treated (BAY–Cyto, blue). (D) A schematic workflow illustrating the employed strategy to identify the inflammatory endothelial cell-state–specific gene modules (endoBBB-M) and evaluation of their relevance in AD brain endothelial modules. (E) Heatmap showing significant enrichment of perfusable BBB-Chip–derived endothelial modules (endoBBB-M1, endoBBB-M3, endoBBB-M7) with AD-associated cortical gene modules (endoM2, endoM4, endo5) described in Figure 1. Color code represents the fold enrichment, with darker blue color indicating higher enrichment. ** FDR < 0.01, *** FDR < 0.001. (F) Density plots of scaled expression of module score showing induction of the endoBBB-M1 module under cytokine treatment compared to control in the BBB-Chip model, with rescue in the BAY-Cyto treatment condition. Kruskal-Wallis test followed by Dunn’s multiple comparisons test. Cyto vs Ctrl: increased expression, adjusted *p*-value < 0.0001, BAY-Cyto vs Cyto: decreased expression, adjusted *p*-value < 0.0001 (also see **Figure S12**). (G) Network plot showing the top 25 hub genes of endoBBB-M1 module. Hub genes include inflammatory mediators such as cytokines and chemokines (IL1B, CXCL5, *CXCL3*), as well as genes involved in TNF (*TNFAIP2*) and NF-κB signaling (*NFKB1*). (H) UMAP (left) and violin (right) plots showing expression distribution of endoBBB-M1 across endothelial states (left), with higher enrichment in the inflammation-associated clusters (Endo-4, Endo-5, Endo-6) (right). (I) Gene ontology enrichment analysis of endoBBB-M1 indicating functions related to lipid storage, NF-κB signaling, inflammatory response, and lipopolysaccharide response. X-axis represents the -log10 (FDR). (J) Cross-validation of endoBBB-M1 relevance across experimental and human AD datasets, showing significant enrichment for DEGs (FDR < 0.05) in cytokine-treated *APOE3* and *APOE4* iBMEC endothelial cells (left panel, also see Figure 3), as well as for genes differentially expressed in AD (middle panel) across multiple human endothelial transcriptomic datasets (Yang et al., 2022; Bryant et al., 2023). Moreover, the endoBBB-M1 module shows significant overlap with genes associated with AD risks (rightmost panel), as identified by GWAS analysis (Grubman et al., 2019). ** *p* < 0.01, *** *p* < 0.001. PFC, prefrontal cortex; EC, entorhinal cortex; ITG, inferior temporal gyrus; V1, primary visual cortex; V2, secondary visual cortex; AD GWAS, Alzheimer’s disease genome-wide association studies.

To systematically capture potential gene-regulatory network associated with this inflammatory iBMEC state (**Figure 6D**), we derived nine endothelial modules (endoBBB-M1 to endoBBB-M9) based on all ECs profiled in the BBB-Chip scRNA-seq and quantified their overlap with the AD-associated brain modules (endoPFC-M2, endoPFC-M4 and endoPFC-M5) (**Figure 6E**). Module overlap analysis demonstrated that endoBBB-M1, endoBBB-M3 and endoBBB-M7 significantly overlapped with the inflammation- and EndMT-associated endoM2 program defined in postmortem human cortical ECs (endoPFC-M2) (**Figure 6E**, **Figure 1**). Among these, we focused on the BBB-Chip-derived module(s) exhibiting opposite expression patterns between cytokine and BAY11-cytokine treatment conditions (**Figure S12B**). Differential module analysis identified endoBBB-M1 as the only BBB-Chip-derived module significantly upregulated in cytokine-treated iBMECs compared to control, with its expression rescued by pre-treatment with the NF-kB inhibitor BAY11-7082 (**Figure S12B, Figure S6F, Table S10**).

Network analysis of endoBBB-M1 hub genes further highlighted central inflammatory mediators including *IL1B*, *TNFAIP2*, *NFkBIA*, *TNFAIP6*, *CXCL1/5/8*, *TNFRSF11B,* and *ICAM1*, together with extracellular matrix–remodeling factors (*MMP10*, *PTX3*) and chemokines (*CCL2*, *CCL20*) (**Figure 6G**). Notably, the module also contained *SVIL*, which we previously found perturbed in AD brains (**Figure 1J–K**). Consistent with these findings, mapping module scores back onto the UMAP confirmed that endoBBB-M1 activity was selectively enriched in the inflammation-associated endothelial state (**Figure 6H**, **Figure 6B**, **Figure S12A**). Gene ontology analysis further revealed strong enrichment for canonical NF-κB signaling, cytokine production, response to lipopolysaccharide, lipid storage, and regulation of inflammatory responses (**Figure 6I**). Importantly, the endoBBB-M1 module significantly overlapped with the cytokine-induced DEGs in *APOE3* and *APOE4* backgrounds from 2D BMECs, with AD-related endothelial signatures in the brain, and with AD-associated risk genes (**Figure 6J**). This demonstrates the robust cross-platform reproducibility of the endoBBB-M1 module, its strong relevance to AD brains, and provides proof-of-concept that inflammation can drive AD-associated endothelial signatures. Together, these results provide strong evidence that inflammatory stimulation drives distinct inflammatory transcriptional state in endothelial cells, marked by activation of a conserved AD-associated inflammatory, EndMT-like gene signatures (endoBBB-M1/endoPFC-M2), which can be effectively suppressed by NF-κB inhibition via BAY11-7082.

## Discussion

The BBB is the critical interface between the brain and periphery where genetic risk factors and environmental stressors can converge to contribute to neurodegeneration. Our data show that cerebrovascular endothelial cells undergo profound transcriptional and phenotypic changes in the context of inflammation and Alzheimer’s disease (AD), displaying EndMT-like signatures marked by cellular elongation, lipid accumulation, and tight junction disruption. Although EndMT has been extensively studied in cancer and cardiovascular disease,^53–57,95,96^ and a recent mouse study suggests it contributes to cerebrovascular dysfunction,^97^ its tie to the cerebrovasculature in the context of dementia in humans has not yet been established. This loss of BBB properties could lead to increased leukocyte infiltration, debris uptake, or pathogen entry into the brain.

Mechanistically, we map this inflammation-driven state to a conserved endothelial program that couples NF-κB signaling to cytoskeletal and junctional remodeling. In the human postmortem cortex, the endoM2 module is enriched for inflammation-related genes and EndMT-associated regulators, and tracks with amyloid/tau pathology, inversely with cognition, and exhibits an *APOE4* dose dependence. We then recapitulate key features *in vitro*: a pro-inflammatory cytokine cocktail induces iBMEC elongation, lipid accumulation, and ZO-1 redistribution, while bulk and single-cell transcriptomics converge on NF-κB–associated reprogramming that overlaps endoM2, including coordinated induction of EndMT drivers (*SNAI1*, *ZEB2*, *SVIL*, *SSH1*) and lipid-storage genes. These shifts parallel those reported in similar contexts: *APOE4* ECs exhibit enhanced uptake of acetylated-LDL (mimicking oxidized LDL) compared with *APOE3*,^98^ TNF-α causes lipid droplet formation in artery and vein ECs^74^ and claudin-5 loss in mouse brain ECs,^99^ and claudin-5 and occludin are lower in AD brain endothelium.^100^ Here, we define the cell state-specific inflammatory transcriptional programs that underlie BBB destabilization and dysfunction. Pre-treatment with the NF-κB inhibitor BAY11-7082 blunts these morphological and transcriptional changes in both 2D iBMECs and a perfusable, multicellular BBB-Chip under flow, indicating that inflammatory signaling is sufficient to impose, and pharmacologic inhibition can attenuate, this AD-like endothelial state. These findings are in line with prior studies demonstrating that ECs show notable interferon signatures in *APOE4* carriers,^101^ and suggest that inflammation, exacerbated by *APOE4* status, can drive vascular dysfunction. Consistent with this, *APOE4* ECs display higher basal inflammatory signatures than *APOE3*, potentially sensitizing them to inflammatory stressors.^102^ This highlights the BBB endothelium as both a site of disruption and a modifiable target in AD pathogenesis.

A strength of this study is that we test the functional relevance of the AD-associated processes in a brain-like BBB environment *in vitro*. The 2D iBMEC model allowed us to rapidly screen different compounds and easily assess key phenotypes (e.g., elongation), while translating our findings into the 3D BBB-Chip model allowed us to verify our findings in a system of increased biological relevance The BBB-Chip transcriptionally captures central human BBB features, with expression of key canonical, junctional, and transporter genes. While cellular elongation is a facile readout in 2D, this particular phenotype is not amenable to 3D monitoring due to the already elongated cellular nuclei in the vasculature at baseline. Instead, we show striking vascular retraction upon cytokine stimulation in the BBB-Chip that is under continuous flow, in addition to lipid accumulation and junctional disruption. These changes were reduced by NF-κB inhibition, supporting a direct mechanistic link between inflammatory signaling and structural barrier dysfunction. Single-cell profiling further revealed discrete inflammatory endothelial state(s) and a BBB-derived gene module (endoBBB-M1) enriched in the inflammatory state, which significantly overlapped with the postmortem human AD-associated endoM2 program and other AD-related signatures, providing compelling proof-of-concept of endothelial vulnerability to inflammation in AD. Thus, the BBB-Chip establishes a flow-capable, cell-resolved platform for mapping disease-relevant states and testing mechanism-anchored interventions.

Another strength is the discovery–replication structure across diverse human snRNA-seq cohorts. We defined endoM2 in a large integrated atlas, validated its enrichment across cortical regions and independent datasets, linked it to aging and *APOE4* gene dosage, and observed selective elevation in clinically diagnosed dementia. We then re-created core features of this program in iPSC-derived BMECs and the BBB-Chip, and showed that pharmacologic suppression of NF-κB signaling reduces both molecular and cellular readouts. Together, these cross-platform replications strengthen the case that a conserved inflammatory/EndMT-like program underlies BBB vulnerability in AD. Our integrated analysis additionally underscores the need for improved methods to isolate, enrich, and sequence endothelial populations from human tissue.^22,39^

Finally, our iPSC-based BBB models demonstrate the feasibility of connecting transcriptional programs to measurable phenotypes and therapeutic rescue. Notably, BAY11 was originally characterized as an NF-κB inhibitor in ECs, where it blocked cytokine-induced adhesion molecule expression and vascular inflammation, providing precedent for its protective role.^79^ While the precise molecular targets of BAY11 are multifaceted, its ability to blunt cytokine-induced vascular remodeling, lipid accumulation, and junctional disruption across models supports the concept of pathway-directed BBB protection. More broadly, these findings establish proof-of-concept that BBB-targeted strategies can potentially mitigate endothelial vulnerability in AD, and position the cerebrovascular endothelium as an accessible, therapeutically tractable interface.

### Limitations of the study

Endothelial nuclei are typically underrepresented on a per-donor basis in standard snRNA-seq experiments; we partially mitigated this limitation by analyzing a large donor cohort and performing cross-validation with datasets specifically enriched for vascular cells. Transcriptomic analyses are cross-sectional in the postmortem brains and thus reflect terminal states, limiting our ability to establish temporality or causality. We focused on cortical ECs and did not explore regional heterogeneity or non-endothelial cell types. Our *in vitro* perturbations model acute rather than chronic inflammatory exposure present in AD, and we note that BAY11-7082 has pleiotropic effects beyond NF-κB inhibition. Future work should incorporate endothelial enrichment protocols and additional brain regions for sequencing analysis,^22,39^ longitudinal or chronic stimulation paradigms *in vitro*, a 3D cellular model that incorporates immune cells and neurons, and more selective or genetic perturbations to clarify mechanisms and therapeutic potential. Ultimately, extending BBB-Chip applications toward modeling vascular resilience and therapeutic rescue in patient-derived settings could establish a roadmap for BBB-targeted interventions in AD.

## Resource Availability

### Lead contacts

Further information and requests for resources and reagents should be directed to the Lead Contacts, Li-Huei Tsai, Md Rezaul Islam, Rebecca L. Pinals.

### Materials availability

New materials generated in this study are available upon request.

### Data and code availability

All data necessary to assess the conclusions of this research are available in the text and supplementary materials. ROSMAP Dataset related to Mathys et al., 2023 were obtained from the authors and are available at Synapse https://www.synapse.org/#!Synapse:syn52293417. Dataset related to Reid et al. 2025 were obtained from the authors and are available at the NCBI Sequence Read Archive under BioProject accession PRJNA1182356. Dataset related to Jeffries et al., 2025 for human aging brains were obtained from https://publications.wenglab.org/SomaMut/. Bulk RNA-seq data from iBMECs and single-cell RNA-seq data will be deposited in the Gene Expression Omnibus. Code will be made available on GitHub upon publication.

## Acknowledgements

R.L. Pinals acknowledges support from the Schmidt Science Fellows program in partnership with the Rhodes Trust and the Burroughs Wellcome Fund Career Award at the Scientific Interface (CASI). M.R. Islam acknowledges support from the EMBO Postdoctoral Fellowship. The authors would like to acknowledge funding through National Institutes of Health grants 3-UG3-NS115064-01S1, 1F32AG072813-01, and 1K01AG083734-01, as well as generous support from the BT Charitable Foundation, Kathleen and Miguel Octavio, the Freedom Together Foundation, The Picower Institute for Learning and Memory, the Robert A. and Renee E. Belfer Family, Eduardo Eurnekian, and David B. Emmes. This work was supported in part by the Koch Institute Support (core) Grant P30-CA014051 from the National Cancer Institute. We thank the Koch Institute’s Robert A. Swanson (1969) Biotechnology Center for technical support, specifically the BioMicro Center. We thank the study participants and staff of the Rush Alzheimer’s Disease Center. Authors thank R.M. Firenze, L. Akay, M. Zoller, M. Wohlwend, D.V. Maydell, and U. Geigenmüller for helpful discussions and feedback. We thank Y. Zhou, T. Garvey, and M. Mazzanti for administrative support. Schematic figures were created in BioRender.

## Author Contributions

Conceptualization: M.R.I., R.L.P., L.H.T.; Methodology: R.L.P. and M.R.I.; Software: M.R.I. and R.L.P.; Formal analysis: M.R.I. and R.L.P.; Investigation: R.L.P., M.R.I., O.K., A.C., E.K., M.N., A.T., M.T.N., A.N., A.J., N.T., E.A., C.F.L.C., C.S.; Resources: M.N., A.E.S., D.A.B., R.L.; Data Curation: M.R.I. and R.L.P.; Writing - Original Draft: R.L.P. and M.R.I.; Writing - Review & Editing: R.L.P., M.R.I., O.K., L.H.T.; Visualization: M.R.I and R.L.P.; Supervision: R.L.P., M.R.I., O.K., and L.H.T.; Project administration: R.L.P., M.R.I., and L.H.T.; Funding acquisition: L.H.T.

## METHODS

## EXPERIMENTAL MODEL AND STUDY PARTICIPANT DETAILS

### Human patient samples from the ROSMAP cohort

Postmortem human prefrontal cortex (PFC; Brodmann area 10) tissue was obtained from participants in the Rush Memory and Aging Project (MAP), a longitudinal, epidemiologic clinical-pathologic cohort study of aging and dementia in older adults from the Chicago area.^103^ Informed consent and Uniform Anatomical Gift Act consent were obtained from all participants and the protocol was approved by the Rush University Medical Center IRB. Participants also provided repository consent allowing data and biospecimen sharing.

We analyzed 10 PFC samples from the MAP study for *APOE3/4* carriers with or without Alzheimer’s disease (AD), classified using the National Institute on Aging (NIA)-Reagan criteria, which integrate Braak stage and CERAD score.^42,43^ AD and control groups did not differ significantly in postmortem interval (< 9 hr) or age at death (86–98 years), as assessed by the Mann-Whitney-Wilcoxon two-sided test. Reported amyloid pathology is the overall amyloid level calculated as the mean of eight brain regions, available for only a subset of the individuals tested. MAP cohort information is summarized in **Table S11**.

### Human iPSC lines and culture

Human iPSC lines used in this study include AG09173 (*APOE3/3* parental from a healthy 75-year-old female donor) and AG09173-1336 (isogenic *APOE4/4* gene-edited); AG10788 (*APOE4/4* parental from a confirmed Alzheimer’s 87-year-old female donor) and AG10788-1207 (isogenic *APOE3/3* gene-edited); and *ZO-1*-mEGFP iPSC line from a healthy 30-year-old male donor (Allen Institute for Cell Science; parental line WTC-11, Coriell GM25256). *APOE3* and *APOE4* cell lines were previously generated by the Picower Institute for Learning and Memory iPSC Facility, with CRISPR/Cas9 genome editing performed as previously described.^104^ iPSCs were cultured on hESC-qualified Matrigel-coated plates (Corning 354277), prepared using 2% v/v Matrigel in DMEM/F12 with HEPES (Gibco) and incubated for 30 min at 37°C. iPSCs were maintained in mTeSR1 medium and passaged using ReLeSR prior to reaching ∼60% confluence (reagents from STEMCELL Technologies).

All iPSC work in this study was conducted under the oversight and approval of MIT’s Institutional Review Board (IRB), known as the Committee on the Use of Humans as Experimental Subjects (COUHES). The research involved only human iPSCs and no human embryos. In accordance with the ISSCR 2021 Guidelines for Stem Cell Research and Clinical Translation, this study falls under Category 1A. All cell materials were obtained, maintained, and used in compliance with institutional biosafety and human subjects policies, as well as the principles outlined in the ISSCR 2021 Guidelines.

## METHOD DETAILS

### Postmortem human brain snRNA-seq datasets, AD-associated gene signatures, and genetic risk genes

Human endothelial single-nucleus RNA sequencing (snRNA-seq) data from the ROSMAP cohorts were obtained and curated from the studies by Mathys et al., 2023 and Reid et al., 2025.^46,47^ In addition, cortical snRNA-seq data related to healthy aging, covering human participants whose ages ranged from 0.4 to 104 years, were retrieved from Jeffries et al., 2025.^52^ All of these snRNA-seq datasets were derived from the prefrontal cortex region of the human brain. Moreover, AD– associated endothelial dysregulated genes were obtained from the studies of Yang et al., 2022 and Bryant et al., 2023.^22,40^ Furthermore, AD-associated risk genes identified through genome-wide association studies (GWAS) were compiled from Grubman et al., 2019 and Yang et al., 2022.

### snRNA-seq analysis of postmortem human brain

#### Human snRNA-seq data processing, integration and annotation

Data analysis was performed using Seurat (v 5.1.0). Data were normalized with NormalizeData, highly variable genes were selected with FindVariableFeatures using top 3000 features, and expression was scaled with ScaleData. Principal component analysis was first performed using the top 30 components, followed by batch correction with Harmony, accounting for both sample and study identifiers as grouping factors. UMAP was computed on the Harmony-reduced space using the first 10 dimensions with parameters n.neighbors = 10, min.dist = 0.2, and spread = 0.1. Neighbor graphs were constructed with FindNeighbors on the same Harmony dimensions, and clusters were identified using FindClusters at the resolution of 0.5. Cluster annotation was guided by vascular zonation markers, including arterial/arteriole genes (*VEGFC*, *ALPL*), venous/venule genes (*IL1R1*, *TSHZ2*), and capillary genes (*MFSD2A*, *SLC7A5*, *INSR*). Clusters were subsequently assigned to major endothelial classes (arterialEC, venousEC, capillaryEC). Cluster-specific markers were identified with FindAllMarkers (only.pos = TRUE, min.pct = 0.25), filtered at FDR < 0.05, and visualized using DoHeatmap.

#### Differentially expressed genes associated with AD pathology and progression

To investigate gene expression changes related to Alzheimer’s disease (AD) pathology and its progression, we applied differential expression analysis using the single-cell framework Nebula (v1.5.3). For pathology-associated analyses, quantitative measures of AD burden were included, such as global pathology score (gpath), neurofibrillary tangle load (nft), and amyloid deposition (amyloid). All statistical models accounted for potential confounders, including donor age, sex, postmortem interval (PMI), study batch, sequencing depth (total reads), and per-cell gene counts. Differentially expressed genes (DEGs) were identified within each endothelial cell type using a false discovery rate (FDR) cutoff of 0.05.

#### Identification of endothelial gene modules in the prefrontal cortex

Genes with zero counts across all endothelial single nuclei were removed. Outlier nuclei were filtered based on detected gene counts using a 3-MAD threshold (log-scale) with scater::perCellQCMetrics. Moreover, genes were retained for downstream analyses if they had more than one count in at least fifty nuclei. Pseudobulk matrices were created using Libra (v 1.7) with min.cells = 50, min_reps =2 and transformed to log2 CPM via edgeR (v 4.2.2), and adjusted for covariates (batch, sex, age at death, PMI) using limma (v 3.58.1). Residuals from this model were used for gene-network construction. Weighted gene co-expression analysis was employed with WGCNA (v 1.72.5) using a signed network with bicor. Soft-threshold powers ranging from 1 to 30 were scanned, and the smallest power achieving a scale-free topology R² ≥ 0.70 was selected. Adjacency, signed TOM, and average-linkage hierarchical clustering were then computed. Dynamic tree cutting was performed with deepSplit = 4, pamStage = TRUE, minModuleSize = 100, and cutHeight = 0.99999. Module eigengenes (MEs) were calculated, and closely related modules were merged at an eigengene dissimilarity threshold of 0.20. Gene-to-module membership (MM) and the corresponding p-values were calculated by correlating individual gene expression profiles with the module eigengenes (MEs). Genes were considered members of a module if their module membership values were greater than 0.30. After applying this threshold, the module eigengenes were recomputed using the filtered gene sets. Signed kME values were used to rank genes within modules, and hub sets were defined as the top 25 genes per module by kME. Module scores for the identified gene modules were computed using the AddModuleScore function in Seurat.

#### Gene ontology, upstream regulators and gene overlap enrichment analyses

Gene ontology analyses for DEGs and gene modules from postmortem brains were performed using either Metascape (https://www.metascape.org/) or Gene Ontology resource (http://geneontology.org). Associations between hub module genes and upstream transcription factors were investigated using ChEA3.^105^ Gene overlap analysis was performed using the GeneOverlap package (v 1.40.0) in R. Gene module score was calculated using the AddModuleScore function in Seurat.

### Immunofluorescence of postmortem human brain

Human brain PFC tissue was prepared as previously described.^106^ Briefly, fresh-frozen tissue was placed on a pre-chilled 10-cm Petri dish on dry ice. A small piece was excised and fixed for 12 hours in 4% (v/v) paraformaldehyde (PFA), washed in PBS, and sectioned into 40 µm slices using a Leica VT1000S vibrating blade microtome. The resulting sections were stored in PBS at 4°C.

To prepare tissues for staining, sections were mounted onto SuperFrost glass slides (Fisher 22-037-246). An ImmEdge pen (VectorLabs H-4000) was used to create a hydrophobic barrier around the mounted tissues. Sections were incubated in a blocking and permeabilization solution (BPS) of 5% (w/v) bovine serum albumin (BSA), 1% (v/v) normal donkey serum (NDS), and 0.3% (v/v) Triton X-100 in PBS for 1 h at room temperature (RT). Sections were then placed in primary antibodies diluted in BPS for 48 hours at 4°C. After three 7-min washes in PBS, sections were incubated in secondary antibodies at a concentration of 1:500 in BPS without Triton X-100 for 2 h at RT. Followed by another three 7-min washes in PBS, sections were then stained with Hoechst (Invitrogen H3570) at a dilution of 1:10,000 in PBS for 1 h at RT. Immunostained sections were once again washed three times at RT in PBS and incubated in 1:20 TrueBlack (Biotium 23007) in 70% ethanol for 2 min. After a final round of 7-min PBS washes, the slides with mounted tissues were dried and coverslipped using Fluoromount-G mounting medium (Southern Biotech 0100-01).

### Confocal microscopy of postmortem human brain

Imaging was performed on a Zeiss LSM900 confocal microscope using ZEN Blue software (Carl Zeiss Microscopy, v3.9). Acquisition settings were maintained uniformly across each image set. Images were acquired at 20× magnification (Zeiss, Objective Plan-Apochromat 20×/0.8). Excitation was performed using 405, 488, 561, and 640 nm lasers on separate imaging tracks. Emission windows were configured with variable secondary dichroics to minimize spectral overlap (410-470 nm, 492-550 nm, 564-625 nm, and 650-700 nm, respectively). Laser-blocking filters were applied to all excitation lines, and a shortpass 470 filter was used with the 405 nm laser. The pinhole diameter was set to one Airy unit (1 AU) for each channel. Laser intensity and detector gain were adjusted to avoid saturation. Z-stacks were set to capture the full sample thickness and step sizes determined by Nyquist sampling.

### Image analysis of postmortem human brain

Confocal image stacks were analyzed in Fiji (ImageJ) with the 3D ImageJ Suite. Channels for PECAM (488 nm Alexa Fluor secondary antibody), supervillin (SVIL) or slingshot protein phosphatase 1 (SSH1) (555 nm Alexa Fluor secondary antibody), and agglutinin (649 nm fluorophore-conjugated primary) were extracted. Vessel structures were defined by summing PECAM and agglutinin channels, thresholding to generate a binary vessel mask, and dilating the binary mask by one voxel across the z-stack. SVIL or SSH1 was thresholded separately to generate a 3D binary image. Fractional coverage was calculated as the proportion of voxels positive for the marker of interest, SVIL or SSH1, within the vessel mask, and mean fluorescence intensity was quantified within the same region of interest (ROI). Quantification was applied uniformly across all samples. Representative images shown were uniformly adjusted by brightness/contrast enhancement and despeckle filtering.

### Generation of PLIN2-RFP reporter iPSC line

The *PLIN2*-RFP reporter iPSC line was generated using the Invitrogen TrueDesign Genome Editor and TrueTag Donor DNA Kit. A construct encoding GSGSGSG (flexible linker)-RFP-P2A-Puromycin was inserted at the C-terminus of the endogenous *PLIN2* gene in AG09173-1336 iPSCs via CRISPR-Cas9 homology-dependent repair. The donor DNA was prepared by PCR amplification and purification of a TrueTag Donor Template (C-RFP-(flox)-Puro) with target homology arms using the following primers: *PLIN2* C1 TRUETAG Fw and *PLIN2* C1 TRUETAG Rv. A 100 μL ribonucleoprotein (RNP) complex was prepared by mixing 1200 ng of TrueGuide sgRNA, 2500 ng of purified donor DNA, and 6250 ng of TrueCut Cas9 protein v2 in Resuspension buffer R. For transfection, 55 μL of the RNP complex was mixed with 55 μL of iPSCs (10,000,000 cells/mL) prepared as a single-cell suspension. The 100 μL mixture was then electroporated using a Neon Transfection system (MPK5000) under the following conditions: 1200 mV for 20 ms with 1 pulse. Following transfection, cells were cultured overnight in mTeSR1 medium containing ROCK inhibitor Y-27632, after which the medium was replaced with fresh mTeSR1 medium. Two days post-transfection, RFP-positive cells were enriched by culturing the cells for 7 days in mTeSR1 medium supplemented with 150 ng/mL Puromycin. Genomic DNA was extracted and purified from the enriched cells using a QIAamp DNA kit. Successful integration of the donor DNA at the end of *PLIN2* Exon 8 was confirmed using TrueTag_C-PuroRFP Verification Fw and Rv primers. Primer sequences are listed below:

PLIN2 C1 TRUETAG Fw: GCCAGGAGACCCAGCGATCTGAGCATAAAACTCATGGAAGTGGCTCAGGTTCTGGA

PLIN2 C1 TURETAG Rv: GTCATCTGTCTGGCCACAGCATGCACTAGTGATAGGGGCAGGTTTACTTGGCCGATCGCATACAGAG

TrueGuide sgRNA: CAUGCACUAGUGAUAGGGGC

TrueTag_C-PuroRFP Verification Fw: GCCCTAGAACCTGGTGCATGAC

TrueTag_C-PuroRFP Verification Rv: GGCTTGCCTTCGCCCTCGGATG

### Differentiation of iBMECs

iBMECs were differentiated similarly to previous protocols, with modifications.^107–109^ iPSCs were dissociated using Accutase, centrifuged (300 rcf, 5 min), and resuspended to single cells in mTeSR1 medium supplemented with 10 µM ROCK inhibitor Y-27632 (reagents from STEMCELL Technologies). iPSCs were plated onto basement membrane-grade Matrigel-coated plates (Corning 354234) at a cell density of 10,000 cells/cm^2^. Culture medium was changed daily throughout both the differentiation and maintenance phases, with 2-3 mL medium volume per 6-well depending on cell density. For days 1-3, the priming medium was DMEM/F12 with GlutaMAX (Gibco), 1% v/v MEM non-essential amino acids (Sigma-Aldrich), and 1% v/v penicillin-streptomycin (Gibco), additionally supplemented with 10 ng/ml BMP-4 (PeproTech) and 6 µM CHIR-99021 (Cayman Chemical). For days 4-7, medium was replaced with endothelial vascular medium (EVM: 2% v/v B27 supplement (Gibco), 1% v/v MEM non-essential amino acids, and 1% v/v penicillin-streptomycin) supplemented with 2 µM forskolin (STEMCELL Technologies) and 50 ng/mL VEGF-165 (PeproTech). On day 6, cells were passaged using Accutase (5 min at 37°C) at a ratio of 1:6 onto collagen I-coated plates (Gibco; 50 µg/mL in DPBS coated on plates for 30-60 min at room temperature). On day 8, cells were selectively passaged using Accutase (30-120 sec at room temperature, with visual confirmation of iBMEC detachment) into endothelial maintenance medium (EVM supplemented with 10 ng/mL VEGF) at a ratio depending on starting confluency of plate (e.g., 1:8 if ∼60% confluent or 1:10 if ∼80% confluent by visual inspection). iBMECs were fed daily with maintenance medium and passaged prior to reaching 80% (typically on day 10), until use on day 12–14. Expression of PECAM (canonical endothelial cell marker) and CD34 (endothelial microvascular marker), along with the absence of EPCAM (epithelial cell marker) and PDGFRb (pericyte cell marker), were evaluated by flow cytometry prior to use (**Figure S6B**).

### Bulk RNA sequencing of iBMECs

On day 12–14 of differentiation, iBMECs were passaged onto collagen I-coated 10cm dishes at a density of 500,000 cells/dish (∼8,820 cells/cm^2^). Culture medium was changed daily (EVM supplemented with 10 ng/mL VEGF) using 10 mL per dish. Four days post-passage, iBMECs were treated with cytokines (TNF-α, IL-1β, and IL-6 each at 1 ng/mL, sourced from PeproTech) vs. control (medium) for 24 hours. Total RNA was isolated using the Direct-zol RNA Microprep kit (Zymo Research) following manufacturer’s instructions. RT-qPCR was performed using 5 ng of total RNA with the RNA to cDNA EcoDry Premix (Takara Bio) and gene expression was quantified from the subsequent cDNA using SYBR green signal from SsoFast EvaGreen Supermix (BioRad). RNA quality was confirmed using the Fragment Analyzer (Agilent Technologies) and RNA-seq libraries were prepared from 100-300ng of total RNA using the NEBNext Poly(A) mRNA magnetic isolation module (New England Biolabs E7490) and NEBNext UltraII RNA library prep kit for Illumina (New England Biolabs E7770). Libraries were prepared following the manufacturer’s recommended protocol using 11-13 cycles of PCR and unique dual indexes. Libraries were validated by sizing using the Fragment Analyzer and quantified by qPCR. Normalized libraries were sequenced on an Illumina NextSeq500 (40nt paired end).

### Bulk RNA sequencing analysis of iBMECs

#### Preprocessing and mapping of samples

Sample preprocessing and downstream analyses of bulk RNA-seq data from iBMECs were performed as previously described.^110^ Base calling and FASTQ file generation were carried out using bcl2fastq (v2.18.0). The quality of raw sequencing reads was assessed with FASTQC (v0.11.5). Alignment of the sequencing reads to the human reference genome (hg38) was performed with the STAR aligner (v2.5.2b). Gene-level count matrices were then produced using the featureCounts function from the Subread package (v1.5.1).

#### Differential gene expression analyses

Genes with raw read counts of at least 5 in a minimum of 50% of the samples were retained for this analysis. Differential expression analysis was carried out using the DESeq2 (v 1.44.0). Genes with adjusted *p*-value < 0.05 after multiple testing corrections were considered as significantly differentially expressed.

#### Gene co-expression analyses

To identify iBMECs-derived modules from the bulk RNA-seq data, genes were filtered to retain those with gene counts (raw) greater than 5 in at least 50% of the samples. Expression values were normalized to library size using the cpm function in edgeR and subsequently transformed as log2(1+CPM). The resulting expression matrix was then transposed to a samples-by-genes format for Weighted Gene Co-expression Network Analysis using WGCNA (v1.61). Sample outliers were identified by computing signed adjacency between samples, estimating whole-network connectivity, and standardizing the connectivity (Z.k). Samples with Z.k values below −3 or with NA values were removed. Network construction was performed by selecting a soft-thresholding power using the pickSoftThreshold function with biweight midcorrelation and a signed network type. Powers from 1 to 30 were assessed, and a power of 7 was chosen to approximate scale-free topology (R² ≥ 0.8). An adjacency matrix was computed using bicor correlations and raised to the soft threshold, followed by conversion into a topological overlap matrix (TOM). The dissimilarity matrix (1−TOM) was used for hierarchical clustering of genes. Modules were identified using the dynamic tree cut method with deepSplit = 3, pamStage enabled, a minimum module size of 50, and a cutHeight of 0.99999. Module eigengenes were computed for each module, eigengene clustering was performed, and close modules were merged with a cutHeight of 0.15. For each gene, module membership (MM) was defined as the correlation between its expression and the module eigengene. Genes with MM ≥ 0.60 were retained as high-confidence module members, and module eigengenes were recomputed based on these filtered genes. To identify hub genes, eigengene connectivity (kME) was calculated for all genes with respect to each module using signedKME from WGCNA. The top 25 genes per module were selected as hubs based on the calculated kME. For network visualization, reduced TOM submatrices were generated for these hub genes, and the strongest 500 edges were retained. Networks were subsequently visualized using igraph (v1.3.1). Functional annotation of modules was performed through Gene Ontology (http://geneontology.org) and Metascape (https://www.metascape.org/). Comparison of the iBMEC modules with AD-brain-derived modules was performed using GeneOverlap package (v 1.40.0).

### *In vitro* phenotypic assays with iBMECs

On day 12–14 of differentiation, iBMECs were passaged onto basement membrane-grade Matrigel-coated (Corning 354234) black-sided glass-bottom 96-well plates (CellVis) at a density of 15,000 cells/well. Culture medium was changed daily using 200 µL per well (EVM supplemented with 10 ng/mL VEGF). Four days post-passage, iBMECs were treated for 24 hours with one of the following conditions: (i) 100 ng/mL lipopolysaccharide (LPS; Sigma-Aldrich), (ii) 1 ng/mL each of cytokines TNF-α, IL-1β, and IL-6 (PeproTech), (iii) 20 µM oleic acid (Sigma-Aldrich), or (iv) control medium. For live-cell imaging of nuclei, lipid droplets, and low-density lipoprotein (LDL) uptake, iBMECs were labeled with Hoechst 33342 (Invitrogen; 1:1000, 30 min) and either BODIPY493/503 (Invitrogen; 1:1000, 30 min) or pHrodo-red LDL (Invitrogen; 1:200, 2 hr; red fluorescence is only visible upon uptake into acidic endolysosomal compartments), followed by a single media wash.

### Live confocal microscopy

Live iBMEC imaging was performed on a Zeiss LSM900 as described for the fixed postmortem human samples, with the addition of a 37°C heated lid and 5% CO_2_ atmosphere. For extended time-course imaging of tight junctions, live iBMECs were imaged on an Agilent BioTek Cytation C10 confocal imaging reader. The imaging chamber was maintained at 37 °C and 5% CO₂. Images were acquired using Gen5 Image Prime software (v3.16) at 20× magnification (Olympus, Plan Fluorite objective) and a 40 µm Nipkow spinning disk. Excitation/emission settings were 472/520 nm for *ZO-1*-mEGFP and z-stacks were set to capture the full cell monolayer thickness. Hoechst staining was omitted due to mild toxicity observed during prolonged imaging. Experiments were conducted with *n* ≥ 3 wells per condition and *n* ≥ 2 regions of interest (ROIs) per well. Representative images are displayed as maximum intensity projections.

### Image analysis for *in vitro* experiments

Nuclei, lipid droplets, and LDL were quantified using custom Fiji (ImageJ) macros (NIH, v2.14.0/1.54g). Threshold values were manually defined for each image batch based on a representative subset of control and treated images, then uniformly applied to generate binary masks. The *Analyze Particles* function was applied, with size constraints set to 10–300 µm² for *APOE4* nuclei, 80–450 µm² for *PLIN2*-RFP-*APOE4* nuclei, and 0.1–12.5 µm² for all lipid droplets and LDL, based on empirical observations from the acquired images, published size ranges, and the resolution limit of the imaging system.^111–117^ Nuclei were quantified in both BODIPY and pHrodo-LDL stained wells (thus doubling the number of data points).

ZO-1 tight junction intensity was quantified using two complementary pipelines. For live-cell imaging of *ZO-1*-mEGFP iBMECs, following thresholding and binarization, images were filtered (3× erode and 3-pixel maximum filters) and bright ZO-1 aggregates that had dissociated from cell borders were identified and removed via ROI-based clearing. Mean fluorescence intensity was then measured. For fixed-cell imaging of anti-ZO-1 stained iBMECs, nuclear masks were subtracted and tight junction structures were detected using the *Ridge Detection* plugin, then integrated intensity was summed across all detected junctional ROIs. SVIL or SSH1 stainings were analyzed as total integrated fluorescence intensity after denoising with the *Remove Outliers* function (radius of 4 pixels).

### *In vitro* drug treatments with iBMECs

Experiments mirrored those of the phenotypic assay described above, with the addition of a one-hour drug pre-treatment prior to the cytokine incubation step. The tested drug panel of previously reported *SVIL/SSH1* inhibitors and related compounds included: 10 µM Sennoside A (Selleck), 10 µM Hypericin (Selleck), 3 µg/mL Wnt antagonist III Box5 (Calbiochem), 25 µM Dvl-PDZ domain inhibitor II (Calbiochem), 10 µM Resatorvid (Selleck), and 10 µM BAY11-7082 (Selleck) (all drugs reconstituted in DMSO).^76–78^ All imaging was performed on the Cytation C10 confocal imaging reader with analysis in Fiji as described above. Experiments were conducted with *n* = 3 wells per condition and *n* = 4 regions of interest (ROIs) per well. Images were analyzed with the same pipeline as described above.

### 3D human iPSC-derived BBB-Chip model

The iPSC-derived BBB in a microfluidic chip (BBB-Chip) was assembled as previously described.^88^ Briefly, BMECs, pericytes, and astrocytes were each individually differentiated from iPSCs, then co-encapsulated at defined cell ratios (5 × 10⁶ cells/mL iBMECs, 0.3125 × 10⁶ cells/mL pericytes, 0.625 × 10⁶ cells/mL astrocytes, 1.25 × 10⁶ cells/mL fibroblasts) within a fibrin-based hydrogel. BBB-Chips were seeded with approximately 3 µL volume in custom-designed microfluidic chips.^88^ Culture medium was changed daily for BBB-Chips, using Endothelial Cell Growth Medium-2 Microvascular (EGM-2MV; Lonza) supplemented with 50 ng/mL VEGF, 10 ng/mL PDGFβ. On day 8-12, BBB-Chips were pre-treated with 5 µM BAY11-7082 (Selleck) for 1 h, then treated with 1 ng/mL each of cytokines TNF-α, IL-1β, and IL-6 (PeproTech) for 18 h. Phenotypic assays and drug screens were performed using live-imaging protocols analogous to those outlined for the 2D monoculture systems and images were analyzed with the pipeline as described above.

### scRNA-seq analysis of BBB-Chip

#### BBB-Chip single-cell sample preparation

On day 10, BBB-Chips were pre-treated with BAY11-7082 (5 µM, 1 h) followed by cytokine stimulation (TNF-α, IL-1β, IL-6; 1 ng/mL each, 18 h). The following day, single-cell suspensions were prepared using the Papain Dissociation System (Worthington Biochemical), with modifications to the manufacturer’s protocol similar to those described for brain organoids.^118^ Briefly, BBB-Chips were washed with DPBS to remove residual media, device ports were scraped to eliminate any adherent 2D cells from seeding the interfaces, and ports were washed again with DPBS. Ports were next filled with collagenase type IV (0.2 mg/mL; Millipore Sigma) and DNase I (2000 U/mL) diluted in Earle’s Balanced Salt Solution with bicarbonate and phenol red (EBSS) and incubated at 37°C, 5% CO₂ for 5 min. BBB-Chips were washed with DPBS then ports were filled with papain solution (20 U/mL papain, 1 mM L-cysteine, 0.5 mM EDTA) and DNase I (2000 U/mL) diluted in EBSS for ∼30 min at 37°C, 5% CO₂ on an orbital shaker (70 rpm). Hydrogels loosened from the chip chambers were transferred into 24-well plates and gently triturated with a P1000 pipette, followed by two additional ∼10 min incubations at 37°C, 5% CO₂ on an orbital shaker (70 rpm), with gentle trituration in between. Once single-cell solutions were visible under a light microscope, digestions were quenched with inhibitor solution (10 mg/mL ovomucoid protease inhibitor with 10 mg/mL BSA) diluted in EBSS, and cells were centrifuged (300 g, 7 min) and resuspended in DPBS containing 0.04% (w/v) BSA. Cell suspensions were filtered and counted, with actual yields ranging from 0.2–0.7 × 10⁶ cells/mL. Live cell suspensions were processed for single-cell RNA sequencing using 3’ GEM-X Universal Multiplexing Chemistry on a ChromiumX (10X Genomics). Libraries were prepared following standard workflows and sequenced on an AVITI24 150nt high output flowcell (Element BioSciences) generating ∼1.6 billion reads across 16 multiplexed samples.

#### Preprocessing, clustering, annotation, and gene module score calculation

Sequencing raw reads were mapped to the human genome (hg38) and the cell-wise gene counts were generated using CellRanger software (v3.0, 10x Genomics). All downstream processing and analyses were performed using Seurat (v 5.1.0). Single-cell RNA-seq count matrices were imported with Read10X_h5 and converted to Seurat objects using CreateSeuratObject. All Seurat objects were merged using JoinLayers and used for downstream analyses. Cells with low quality were filtered out from the downstream analyses. For each cell, the mitochondrial transcript fraction was calculated using the PercentageFeatureSet function in Seurat, and the distribution across cells was visualized with violin plots. In addition, we examined two standard quality control metrics: the total number of transcripts detected per cell and the number of unique genes detected per cell. Cells were retained if they contained 500–7,500 detected genes. Genes detected in fewer than three cells were removed from downstream analyses. The dataset was subsequently normalized using NormalizeData, highly variable top 3000 genes were identified with FindVariableFeatures, and expression values were scaled using ScaleData. Next, dimensionality reduction was performed with principal component analysis on the top 30 principal components using RunPCA and data were harmonized across samples with Harmony (v 1.2.0). Harmonized data were then used to compute nonlinear embedding with RunUMAP using spread = 5 and min.dist = 0.6. A neighbor graph was later constructed with FindNeighbors, and clustering was performed with FindClusters at resolution 0.2. Major cell classes were annotated by combining canonical marker inspection, de novo marker discovery, and external reference enrichments. Canonical vascular markers were visualized with FeaturePlot. Cluster specific markers were identified with FindAllMarkers (only.pos = TRUE, min.pct = 0.25); significant genes were retained at FDR < 0.05. To further support endothelial annotations, curated endothelial gene panels from prior studies^22,119^ were curated and their expression patterns were displayed as dot plots. Final major labels (e.g., Endothelial cells, Pericytes, Fibroblast, Astrocytes) were assigned by integrating these lines of evidence. Gene module scoring for selected pathways or other supervised analyses (e.g., lipid storage and endoM2 associated BAY-Cyto-rescued DEGs) was performed using AddModuleScore function in Seurat. To identify endothelial heterogeneity in the BBB-Chip model, endothelial cells were further processed and reanalyzed as described above. The data were normalized and harmonized across samples, followed by dimensionality reduction using principal component analysis (PCA). The top 30 principal components were used for clustering at a resolution of 0.3 to define distinct endothelial cell states, which were subsequently visualized using UMAP plots. We applied FindAllMarkers (only.pos = TRUE, min.pct = 0.25) in Seurat to detect endothelial cell state–specific marker genes as described above. Differentially expressed marker genes with a false discovery rate (FDR) < 0.05 were considered significant and used for gene ontology analyses.

#### Differential gene expression and weighted gene co-expression analyses in BBB-Chip scRNA-seq data

Differentially expressed genes (DEGs) among treatment conditions in the BBB-chip were determined using MAST (v 1.20.0). Significant genes were defined as those with FDR < 0.05 and absolute avg_log2FC ≥ 0.1. Weighted gene co-expression network analysis in endothelial cells from BBB-chip was performed as previously described.^120–122^ Briefly, “metacells” were generated by averaging gene expression across each cell and its 25 nearest neighbors using a k-nearest neighbors approach. Metacells were normalized, scaled, and reduced by principal component analysis (PCA) and data were harmonized across samples. A data-driven soft-thresholding power was selected using TestSoftPowers and PlotSoftPowers, choosing the first power at which the scale-free topology model fit (SFT R²) exceeded 0.8. The selected soft power was applied to construct a signed co-expression network, and the topological overlap matrix (TOM) was computed to quantify network connectivity. Gene modules were identified using a minimum module size of 25. Each module was summarized by its eigengene, defined as the first principal component of the module’s gene expression matrix, and eigengenes were used to compare between experimental groups. Gene membership within modules was assessed by correlating individual gene expression with the corresponding eigengene. Hub genes were identified based on intra-modular connectivity (kME), and the top hub genes per module were visualized using the igraph package (v1.3.1). Gene ontology analyses for DEGs and modules from the BBB-chip were performed using either Gene Ontology resource (http://geneontology.org) and Metascape (https://www.metascape.org/). Comparison of the BBB-Chip modules with AD-brain-derived modules, AD-associated endothelial gene signatures and risk genes, was performed using GeneOverlap package (v 1.40.0).

### Immunofluorescence of iBMECs and BBB-Chips

After live-cell imaging, 2D monolayers (3D co-cultures) were briefly washed with PBS, fixed with 0.1% (v/v) glutaraldehyde, 2% (v/v) paraformaldehyde (PFA) in PBS (4% (v/v) PFA in PBS) for 20 min (30 min), washed three times with PBS over 30–60 min, then permeabilized with 0.1% v/v (0.5% v/v) Triton-X for 20 min (1 h). All steps were performed at room temperature on an orbital shaker unless otherwise noted. Plates were then washed three times with PBS for 5 min (20 min) each, incubated in blocking buffer (2% w/v BSA, 3% v/v Normal Donkey Serum in PBS) for 30 min (2 h), then incubated in primary antibodies overnight for 16-18 h at 4°C. The next day, plates were washed three times with PBS over 30–60 min and incubated with respective secondary antibodies for 1 h (2 h). Plates were then washed three times with PBS over 30–60 min and stored at 4°C until imaging.

## QUANTIFICATION AND STATISTICAL ANALYSIS

For imaging experiments, quantification was performed in Fiji as described in the Methods section and Python was used for statistical analysis. For sequencing data, statistical analyses were performed in R (v 4.1.2, v 4.4.0) and Prism (v 8.3.1). Statistical details of experiments and sequencing data analyses can be found in figure legends and the Methods section.

